# Scope for sympatric giant-dwarf speciation driven by cannibalism in South-American annual killifish (*Austrolebias*)

**DOI:** 10.1101/121806

**Authors:** Tom JM Van Dooren, Henri A Thomassen, Femmie Smit, Andrew J Helmstetter, Vincent Savolainen

## Abstract

A trophic radiation in the South-American annual killifish genus *Austrolebias* has led to the evolution of large specialized piscivores from small generalized carnivores. It has been proposed that this occurred in a single series of vicariant speciation events. An alternative hypothesis is denoted giant-dwarf speciation: piscivores would have evolved in sympatry by character displacement and cannibalism. We test the plausibility of both scenarios using size measures combined with distributional data and new phylogenetic trees based on mitochondrial and nuclear molecular markers.

Our analysis uses historical biogeography models and Ornstein-Uehlenbeck processes describing trait evolution across the posterior distributions of phylogenetic trees. Large species most likely evolved three times from small ones. For the clade containing *A. elongatus*, we argue that vicariance was not involved in the origin of these large and specialized piscivores. They experience stabilizing selection with an optimum shifted towards larger bodies and longer jaws. The branch leading to this clade has the fastest evolving jaw lengths across the phylogeny, in agreement with expectations for giant-dwarf speciation. For *A. wolterstorffi*, the support for giant-dwarf speciation is weaker. When the species is placed at the root of *Austrolebias*, ancestral reconstructions are unreliable and vicariance cannot be ruled out. For the remaining large species, we can reject vicariance and giant dwarf speciation. Our results give rise to two new additional scenarios for the evolution of specialized piscivores. In the first, two successive speciation events in sympatry or parapatry produced large and piscivorous species. In the second, the immigration of a different annual killifish genus (*Cynopoecilus*) in the Patos area of endemism has contributed to in-situ diversification of *Austrolebias* species.

## Introduction

Within the annual killifish genus *Austrolebias*, large piscivorous species evolved from small ancestors in a trophic radiation (Costa 2009, 2011, Figure one). According to Costa (2009, 2011), this occurred in a single series of successive vicariant speciation events with gradual evolution of specialized piscivory. This *gradual allopatric* hypothesis involves the least amount of ecological interactions among the emerging *Austrolebias* species, therefore other *Austrolebias* species already present or fish from different genera became the prey. A very different hypothesis is based on the giant-dwarf eco-evolutionary scenario proposed by Claessen *et al* (2000) and Dercole and Rinaldi (2002) and we therefore call it *giant-dwarf* speciation. First, coexisting dwarf and giant morphs emerge within populations as a result of size-dependent cannibalism and plastic growth rate variation (Claessen *et al* 2000). When intraspecific cannibalism subsequently evolves into piscivory of a smaller species, a sympatric species pair of piscivore (giant) and prey (dwarf) can be the outcome (Dercole and Rinaldi 2002). The possibility of speciation driven by cannibalism has been little investigated (Nosil 2012).

The hypotheses proposing either gradual evolution in a single series of vicariant speciation events or giant-dwarf speciation involve contrasting patterns of the spatial distributions of ancestral and descendant species and of trait divergence between species. This offers several possibilities to falsify these hypotheses and make progress in our understanding of speciation in *Austrolebias*. The *gradual allopatric* hypothesis will be rejected when the probability that vicariance occurred can be fixed at zero in the biogeographic model (Ree *et al* 2005, Ree and Smith 2008, Matzke 2014) or when it occurs with low probability at nodes where large species evolved. The *giant-dwarf* scenario will be rejected by a biogeographic analysis when that is the case for the probability of sympatric speciation. Phylogenetic comparative methods (e.g., Ingram and Mahler 2014) will reject the *gradual allopatric* scenario if we can demonstrate that large species or specialized piscivores either originated multiple times or when selection regime shifts occur for both body size and jaw length on the same branch. If giant and dwarf species emerge in a sympatric process of evolutionary branching by negative frequency-dependent selection (Geritz *et al* 1998), disruptive selection causes large phenotypic differences to emerge from an ancestral population with much less phenotypic variation. Since a minimum size difference is often necessary for cannibalism to be possible (e.g., Persson *et al* 2000), one can assume that a relatively rapid and large size and gape change is necessary for diversification by cannibalism. The *giant-dwarf* hypothesis will be rejected when phylogenetic comparative methods assign low probabilities to the occurrence of a selection regime shift on branches where a large species evolved from small and when a potential *giant-dwarf* speciation did not occur with relatively fast trait changes.

We briefly summarize the current state of knowledge on species coexistence and phylogeny of *Austrolebias.* About 45 species are recognized within the South-American annual killifish genus *Austrolebias* (Costa 2006) and new species are being described regularly. *Austrolebias* species mostly occur in temperate zones. They live in temporary ponds, where seasonal drought kills all adults and juveniles present. Populations persist by means of egg banks in the soil containing diapausing embryos. When rains refill the ponds, alevins hatch from the egg bank. In these ephemeral environments, annual killifish are very often the only fish present. However, temporary ponds can become connected to more permanent water bodies and non-annual fish can then invade until the next drought removes them. Coexistence of different annual fish species is very common in ponds where *Austrolebias* occur. From hatching on, most species likely interact with other annual fish and much less so with non-annuals.

Piscivory is uncontested for the largest species such as *A*. *elongatus*: entire fish have been found in their guts (Costa 2009). Using a phylogenetic tree based on morphological characters, Costa (2009, 2011) found these piscivores in an apex position in the cladogram and concluded that large sizes, sporadic and specialized piscivory evolved gradually within the genus in a sequence of vicariant speciation events. A biogeographic reconstruction of species ranges across large areas of endemism (Costa 2010) found many cladogenetic events in the genus without vicariance nor dispersal. Whether all speciation events are allopatric was not tested using statistical inference. Costa (2010) also hypothesized that *Cynolebias,* another genus with large species, is the sister genus of *Austrolebias*. Large *Austrolebias* generally coexist with small *Austrolebias* and in some regions with annual *Cynopoecilus* species (Costa 2016) or with annual species from other genera (Chaco region). *Cynolebias* species usually coexist with small *Hypsolebias* (Costa 2010, Costa *et al* 2018). A recent biogeographic reconstruction of the annual killifish genus *Cynopoecilus* (Costa 2016) inferred that speciation events within the genus were associated with southward migration into the Patos area of endemism. It could be that the presence of annual fish from genera different from *Austrolebias* has provoked the evolution of large piscivores, for example when these became a prey species. A test of *giant-dwarf* speciation should therefore investigate whether piscivory evolved in the Patos or Chaco areas where coexistence with other genera occurs.

Phylogenetic trees of *Austrolebias* based on mitochondrial DNA sequences (mtDNA, Garcia *et al* 2000, 2002, 2014) suggest repeated evolution of large species and therefore seem to reject the gradual allopatric scenario. However, these trees had insufficient support at relevant nodes in the trees to falsify Costa’s (2010) hypothesis. A simple biogeographic reconstruction involving no modelling and not taking phylogenetic uncertainty into account (Garcia *et al* 2000) assigned two origins of large species to sympatric speciation events.

We will test the support for Costa’s (2011) *gradual allopatric* scenario for the evolution of large piscivores and for *giant-dwarf speciation*, using (1) two sets of posterior distributions of molecular phylogenetic gene trees, one obtained by modelling nuclear ribosomal DNA and one based on three mitochondrial markers (mtDNA); (2) a phylogenetic comparative analysis of trait variation assessing selection regime shifts and rates of trait change and (3) a parametric biogeographic analysis. Our analysis will take phylogenetic uncertainty into account throughout.

## Materials and Methods

We used individuals from laboratory populations in the Animal Ecology Lab in Leiden, the Netherlands (Supplementary Table S1) complemented with samples of *Austrolebias monstrosus* and *vandenbergi* from the field. Lab populations were obtained from field trips or expert breeders. They are systematically maintained by crossing individuals from the same clutch, therefore individuals from the same population of origin are kin. The sample consisted of 112 individuals of species in the *Austrolebias, Hypsolebias*, *Ophthalmolebias*, *Nematolebias* and *Spectrolebias* genera (Costa 2010). We added *Cynolebias albipunctatus* to confirm the monophyly of *Austrolebias* and to determine whether *Cynolebias* is its sister taxon. Individuals of *Aphyolebias schleseri* (Costa 2003), *Pterolebias longipinnis* (Garman 1895) and *Leptolebias citrinnipinnis* (Costa *et al* 1988) are included as more distant outgroups. Among the 22 *Austrolebias* species included in our analysis, eight species are commonly denoted as large and form a single monophyletic group in Costa (2006, 2010). These are *A. vazferreirai, cinereus, robustus, wolterstorffi, monstrosus, elongatus, prognathus* and *cheradophilus*. There is a single large species missing from our analysis (*A. nonoiulensis*), which is similar to *A. robustus*, *cinereus* and *vazferreirai* (Costa 2006). We were unable to obtain recent samples from it.

### DNA extraction and amplification

In the African annual *Nothobranchius* killifish, phylogenetic trees using nuclear markers had poor resolution at the gene level (Dorn *et al* 2011, 2014). Therefore concatenated sequences (Dorn *et al* 2014) or a multispecies coalescent were used to obtain a species tree where half of the nodes had nodal support above 80% posterior probability (Dorn *et al* 2011). Remarkably, a study on South-American annual *Simpsonichthys* subgenera using 842 bp of mtDNA markers (Ponzetto *et al* 2016) obtained nodal support values above 80% for most nodes at the interspecific level suggesting that the lack of support in previous trees for *Austrolebias* could be due to limited sequence length. We decided to investigate a minimal combination of individuals, taxa and sequences that could be just considered appropriate (Figure 1 in Philippe et *al* 2017). We opted for a small number of nuclear and mitochondrial sequences of most large *Austrolebias* species and a set of small ones sufficiently covering several biogeographical areas and with several individuals per species when possible.

**Figure 1.**
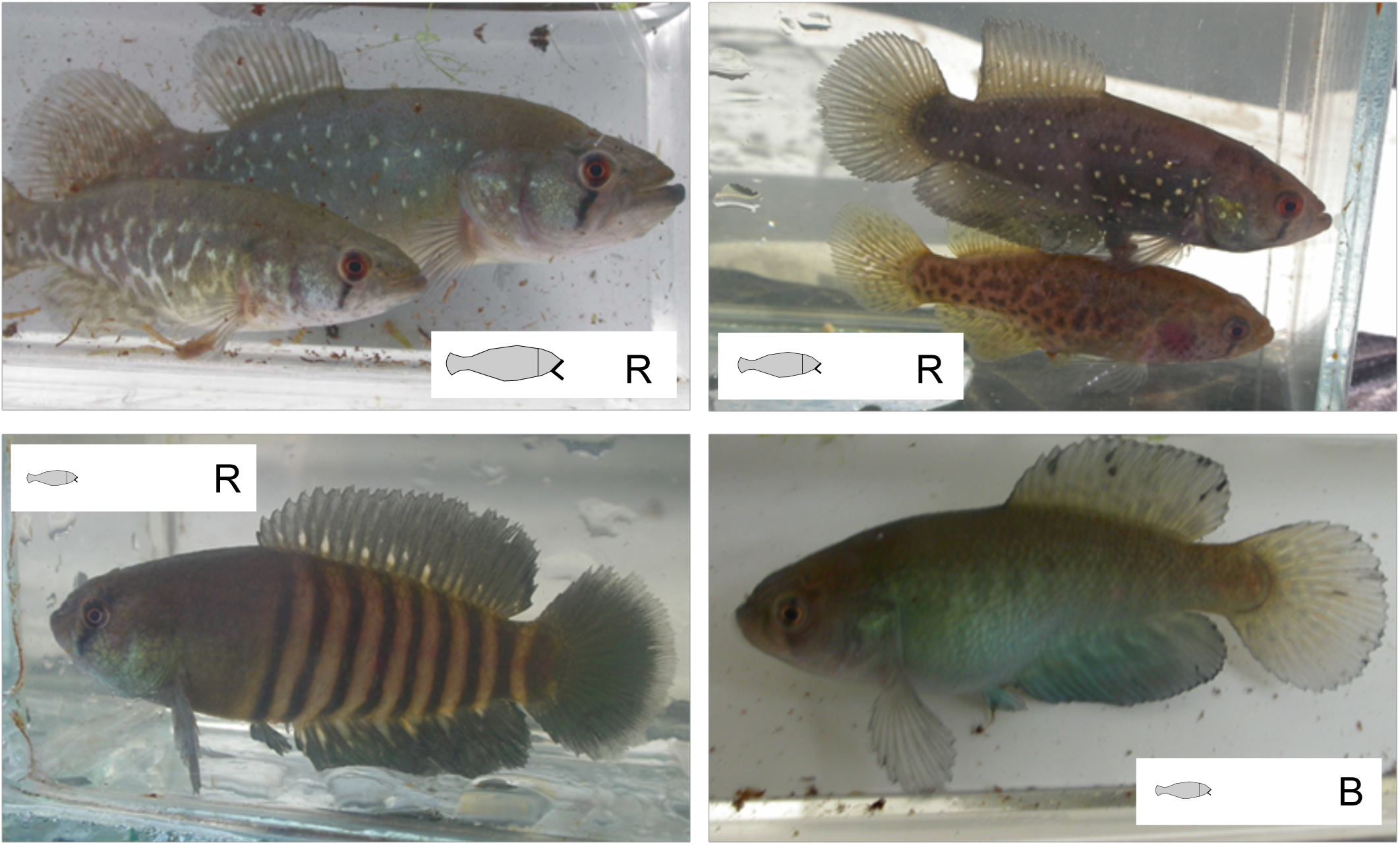
Photos of different *Austrolebias* species. Top left: *A. prognathus* female (front) and male (large species, “Salamanca” population of origin); Top right: *A. wolterstorffi* female (bottom) and male (large species “Canal Andreoni” population); Bottom left: *A. luteoflammulatus* male (small species, “La Pedrera”); Bottom right: *A. cinereus* male (large species, “Arroyo de las Viboras”). The insets in each photo refer to the trait representations for body length and relative jaw length and to symbols for areas of endemism used in Figures two, three and five.

DNA was extracted from fin clips, muscle and liver tissue using the Qiagen DNEasy Blood and Tissue kit following the manufacturer’s protocol and used for direct amplification by Polymerase Chain Reaction (PCR) of mitochondrial sequences of cytochrome-*b* (*cytB*), 12S ribosomal DNA (rDNA), 16S rDNA and nuclear 28S rDNA (3 regions). Table S2 lists primers used for PCR and sequencing and Table S3 PCR reaction conditions. PCR products were cleaned using the Promega Wizard SV Gel and PCR Clean-Up System (Promega, Madison, Wisconsin, USA) and sent to a commercial sequencing facility (Macrogen Inc., Korea, www.macrogen.com).

Sequencing reactions were carried out using our supplied primers and the sequence products were run on an ABI3730XL genetic analyser.

### Sequence analysis

Electropherograms were edited in Sequencher 4.1.4 (GeneCodes, Madison, Wisconsin), and sequences aligned using ClustalX 1.83 (Thompson *et al* 1997; Jeanmougin *et al* 1998) using default parameters (gap opening = 10.00, gap extension = 0.20, delay divergent sequences = 30%, DNA transition weight = 0.50). Alignments were straightforward for cytB, but contained indels in the rDNA sequences. We identified stems and loops in the mitochondrial rDNA sequences based on an alignment in RNAalifold (Bernhart *et al* 2008) resulting in improved alignments containing few ambiguous bases except for some regions within indels. Ambiguous regions were removed from the alignments before further analysis of the 12S and 16S sequences. Indels were coded as presence-absence using simple indel coding in SeqState 1.37 (Müller 2005). The final alignments used for the phylogenetic analyses contained 1590 (mitochondrial) and 727 (nuclear) base pairs and an additional 82 (mtDNA) and 27 (28S rDNA) binary indel-coding characters.

### Mitochondrial and nuclear gene trees

Bayesian phylogenetic inference was carried out in MrBayes 3.2.5 (Huelsenbeck and Ronquist 2001; http://mrbayes.csit.fsu.edu). Datasets for the mitochondrial and nuclear markers were analysed separately but with each subset concatenated. The mtDNA sequences are jointly inherited. The nuclear sequences are segments of a single gene and it is unlikely that introgression would disrupt them. Each dataset was partitioned into separate character sets for the DNA sequence and indels per marker. Character sets for 12S and 16S mtDNA sequences were further partitioned into stems and loops. We fitted different substitution models to determine which one fitted each dataset best, with parameters shared between partitions or not and estimated marginal likelihoods of each model using the stepping stone approximation (Xie *et al* 2011). Based on these comparisons, General Time Reversible models with a proportion of invariant sites and gamma distributed rate variation (GTR+I+G) were implemented, with different parameter estimates per partition of DNA sequence data, except for the stems in the 12S and 16S alignments, where we used the doublet model (Schöniger and von Haeseler 1994). Changes in indel state were fitted with binary models. A doublet model would also be appropriate for 28S stems. We aligned our partial 28S rDNA sequences to the best-matching complete sequence of the gene we could find (*Oreochromis auratus*), for which we determined its secondary structure using RNAfold (Hofacker *et al* 1994). Assuming that our aligned sequences would have stems and loops as in the complete sequence, we observed that stem distributions differed when the structure was determined using minimal free energy or ensemble prediction, that stems often consisted of a sequence in our data and another one not in it, and that some stems consisted of a pair in two different partial 28S sequences. This would cripple an implementation of the doublet model for 28S stems and we used the GTR+I+G model instead.

Per run, base frequency parameters were estimated assuming a Dirichlet distribution. We used default prior settings in MrBayes 3.2.5 except for the prior on branch length (exponential with parameter 20). Per model, two separate runs of four Markov Chain Monte Carlo (MCMC) chains were run until convergence to a stationary regime allowed sampling at least 4000 trees spaced by 5000 generations. The selection of phylogenetic hypotheses continued on the basis of 50% majority consensus trees for these samples, by removing individuals from the dataset and refitting the model. Our sample contained 24 pairs or groups of sibs (same species/population).

We preferred consensus trees with each sib group as sister branches and removed some individuals to achieve such a configuration (single individuals not grouping with several others, individuals with missing data). We checked for long-branch attraction by removing individuals with long branches (Bergsten 2005), and removed subsets of species from the dataset when long-branch attraction was identified. We also ran models for data subsets with only those individuals with data for all mitochondrial or nuclear markers.

### Species trees

The biogeographic and phenotypic analyses below require species trees. These are referred to as the true bifurcation history of species (Mallet *et al* 2015), when based on many sequences.

Nearly all comparative methods and biogeographic reconstructions assume phylogenetic trees to represent the true bifurcation history. All discordance between gene trees is regularly assumed to be due to lineage sorting (Liu and Pearl 2007). This assumption facilitates phylogenetic tree reconstructions by ignoring part of the true history, namely introgression and hybridization between species (Wallis *et al* 2017).

Lab data demonstrate that at least some species pairs in *Austrolebias* can be hybridized (Oviedo *et al* 2016). It is therefore reasonable not to *a priori* consider just a single species tree based on all markers concatenated or on coalescent methods, or to try to obtain a species phylogenetic network using next-generation sequencing for which methods to analyse phenotypic divergence and to fit biogeographical models are limiting. In this first assessment of hypotheses regarding size divergence in *Austrolebias* using molecular phylogenetic trees, we constructed two alternative gene bifurcation histories, one for the concatenated mitochondrial sequences and a second for the 28S sequences. We investigated an additional set of nuclear DNA markers, but did not obtain separate gene trees with sufficient support for further similar analysis. Analyses based on the multispecies coalescent or concatenation of nuclear markers in different genes will be presented elsewhere.

We constructed species trees from mtDNA and 28S gene trees separately using the GLASS approach (Mossel and Roch 2010; Liu *et al* 2010). Gene trees were forced to be ultrametric using the penalized likelihood method of Sanderson (2002) with parameter lambda fixed at value zero (variable rates). Morphological and biogeographical analyses focused on the individuals of the genus *Austrolebias* and we pruned the trees accordingly.

For illustration and to understand potential pitfalls in the analysis, we first carried out phenotypic and biogeographic modelling on the mtDNA and 28S consensus trees. This was then repeated on random samples of 500 trees from the posterior distributions and results were summarized through averaging parameter estimates and predictions across the samples. A first test of the *gradual allopatric* hypothesis consisted in inspecting the posterior distribution for the number of times large species evolved within the genus *Austrolebias*.

### Phenotypic trait analysis

We analysed morphometric summary statistics of field samples obtained from the tables of Costa’s (2006) revision of the *Austrolebias* genus. Maximum and minimum values were reported per trait making allometry analyses requiring individual data or calculation of size-independent principal components impossible. We did not use the minimum standard lengths reported. Fish have indeterminate growth and information on individual age is lacking. Traits pertaining to our hypothesis are sex-specific maximum standard lengths and maximum and minimum lower jaw lengths given as a fraction of head length. Prior to further analysis, we calculated scores for the most important principal components (PC) of (1) the two size traits and (2) the four measures of jaw length. The scores were used as body size and relative jaw size in further analysis. We did not carry out phylogenetic PC’s that are fixing trait correlations between species. These correlations are derived from a specific model for trait evolution, complicating further analysis of the PC’s that are calculated (Uyeda et al. 2015). We did not *a priori* assign eight species as large in our comparative analysis of trait evolution. We fitted Ornstein-Uhlenbeck (OU) models with a single or multiple optima for stabilizing selection (Butler and King 2004) to species trait values and each phylogenetic tree. Evolutionary OU models for continuous trait values assume that trait changes along branches of a phylogenetic tree occur according a process where the species trait means are attracted to one or several central values (optimum values), while species traits are additionally changed by a Brownian motion process. For each species tree sampled from the posterior distributions of gene trees, we determined the number of selection regimes and shifts towards large species in the OU model by applying an automated routine with a modified AICc to the size and jaw scores combined (surface R library; Ingram and Mahler 2013). Only the 500 AICc selected models are included in the averaging of parameters and predictions. To calculate average differences between estimated selection optima, we carried out a parametric bootstrap on each selected model and averaged across the replicates and trees. We limited bootstrap replication to ten per tree, because trees recur in the samples.

The automated surface procedure can lead to overfitting (Ho and Ané 2014) and surface and its alternative l1ou (Khabbazian et al. 2016) both assume that traits are independent, conditional on the shifts. To assess whether observed AICc decreases were sufficient to prefer such OU models with selection regime changes, we carried out simulations. For a sample of 100 trees from each posterior, we fitted bivariate Brownian motion (BM) and OU models with a single optimum to the data, allowing for correlations between traits. On the basis of each fit, 100 new datasets were simulated and the surface procedure repeated. This resulted in a distribution of AICc changes in cases where the BM or OU model is true, and the quantiles and tail probabilities of this distribution allow us to assess whether the BM or OU model can be rejected and a model with selection regime changes preferred for that tree.

Evolutionary models fitted to phenotypic and phylogenetic data should take measurement error into account (Silvestro *et al* 2015). We carried out an approach similar to SIMEX extrapolation (Cook and Stefanski 1994) to assess potential effects of measurement error. Pseudo data sets were generated by adding random errors of increasing variance to the length and jaw scores and we ran the automated model selection procedure on each pseudo data set using a random tree from the posterior. Low sensitivity of the results to increasing values of the added measurement error would suggest that measurement error in our data has limited effects on inference.

To investigate relative rates of trait change, we fitted stable distributions as the model for trait changes (Elliot and Mooers 2014) to each of the species trees in the samples and the size and jaw scores separately. These distributions can be heavy-tailed and thus enriched for large changes. In the results, we inspected whether relatively large changes occurred on branches towards large species by calculating ranks of the estimated median rates of trait change on each branch. The averages of these ranks for particular branches are reported.

### Biogeographical analysis

Four areas of endemism occur among the species in this analysis (Costa 2010, Figure two). We used these areas as discrete categorical species ranges. Each extant species occurs in a single area, but the analysis allows for multi-area ranges in ancestors. Two areas in which small *Austrolebias* occur are lacking in our data, namely the Iguaçu River basin inhabited by *A. carvalhoi* and *araucarianus* (Costa 2006, 2014) and the upper Uruguay River basin with *A. varzeae* (Costa 2006).

To reconstruct ancestral ranges at particular nodes, we used an extension of the likelihood framework and modelling approach of Ree (Ree *et al* 2005, Ree and Smith 2008) as implemented in the BioGeoBEARS package (Matzke 2014). It estimates anagenetic migration and extinction along branches and different cladogenetic scenarios at each node: a jump dispersal to a different area in one descendant, vicariance of a multi-area range, sympatric speciation within a single-area range, and subset speciation when a multi-area range occupied by the ancestor remains occupied by one descendant and the other descendant becomes limited to a single area. Weaknesses pointed out by Losos and Glor (2003) are reduced in newer methods that allow for range changes in between speciation events but these are often inferred to be absent when jump dispersal is allowed (Ree and Sanmartin 2018). In order not to *a priori* bias the overall plausibility of certain cladogenetic scenarios, we first implemented a model with the largest possible number of parameters that could be estimated and applied it to the samples of species trees. Simplified models without vicariance or without sympatric and subset speciation were fitted too and the average change in AICc reported. To assess the probability of vicariance and sympatric speciation at specific nodes in the full model, we used stochastic mapping (100 replications) given the tip data and model parameter estimates (Matzke 2014) and averaged probabilities of events across the species trees.

## Results

### Mitochondrial and nuclear gene trees

In the analyses of the mitochondrial data, 5 or 7 individuals were removed from the dataset when the model was fitted to individuals with full or partial data, respectively, because they did not group with kin or with all other individuals from the same species (Supplementary Material Figures S4-S6). The resulting phylogenetic hypothesis for the mitochondrial markers resolved all species (consensus species tree Figure 3, gene tree Fig. S6) and grouped the large *Austrolebias* species in three well delineated and separated clades: only 4% of the trees in the posterior included two clades with large species. Overall, partition posterior probabilities are above 0.8 for 80% of the nodes within the genus *Austrolebias*, when within-species nodes are excluded. The consensus gene tree for the nuclear 28S data showed long branch attraction between *A. vandenbergi* and *Pterolebias* and *Aphyolebias* outgroups (Figures S1-2). We therefore removed these outgroups. Additionally, five more individuals were removed that did not group with kin or conspecifics (Fig. S3). For every tree in the posterior, we obtained a phylogenetic hypothesis with the large species in three clades. These are again well separated, but *A. robustus* is paraphyletic to *A. vandenbergi*. The mitochondrial and nuclear phylogenetic trees reject Costa’s (2010) hypothesis with a single origin of large *Austrolebias*.

**Figure 2.**
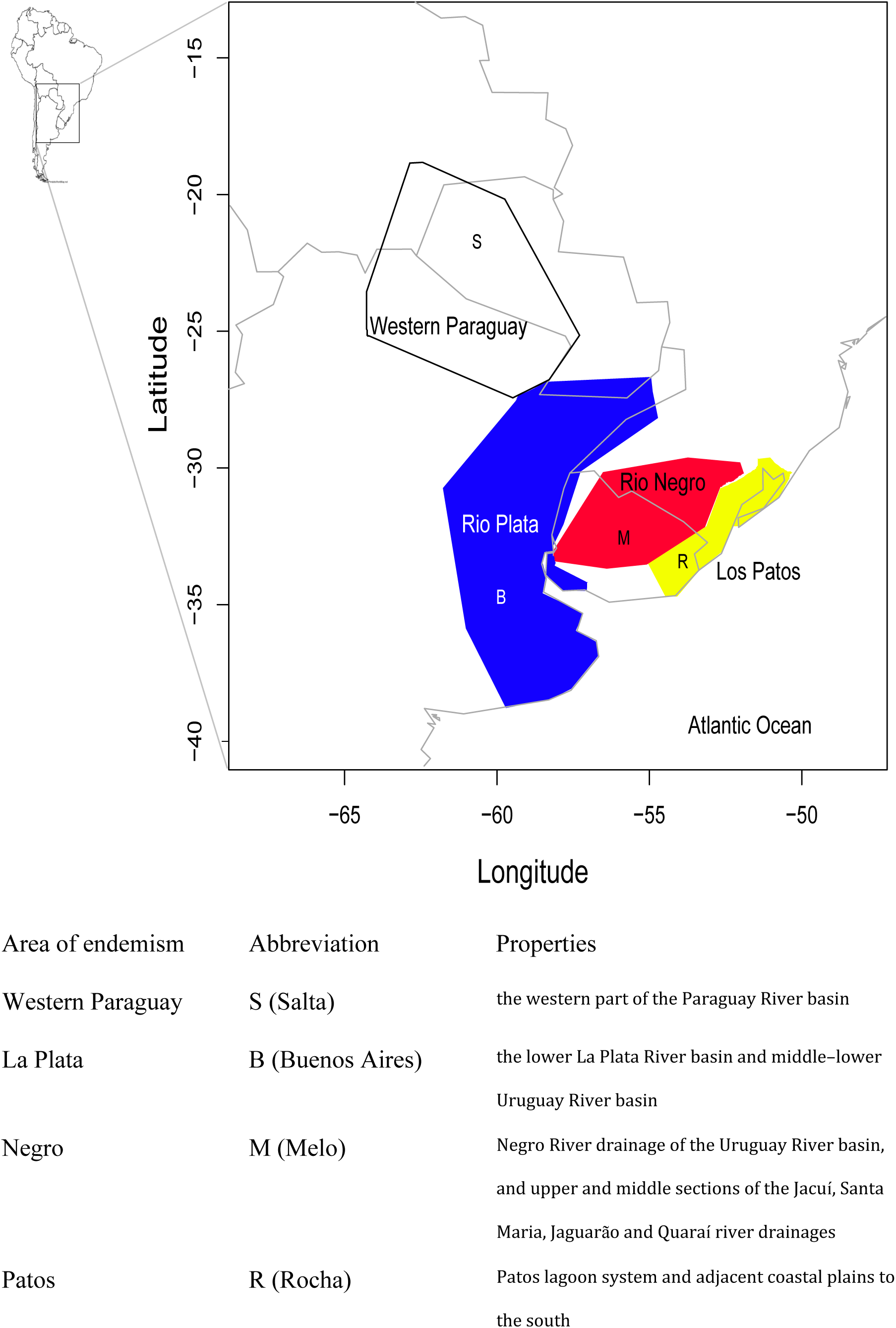
Areas of endemism. The geographic areas used for analysis are indicated on a map of South-America. The letters used to represent them in Figures one and five are given, together with a brief description of each area. All *Austrolebias* species in the analysis occur in a single area.

**Figure 3.**
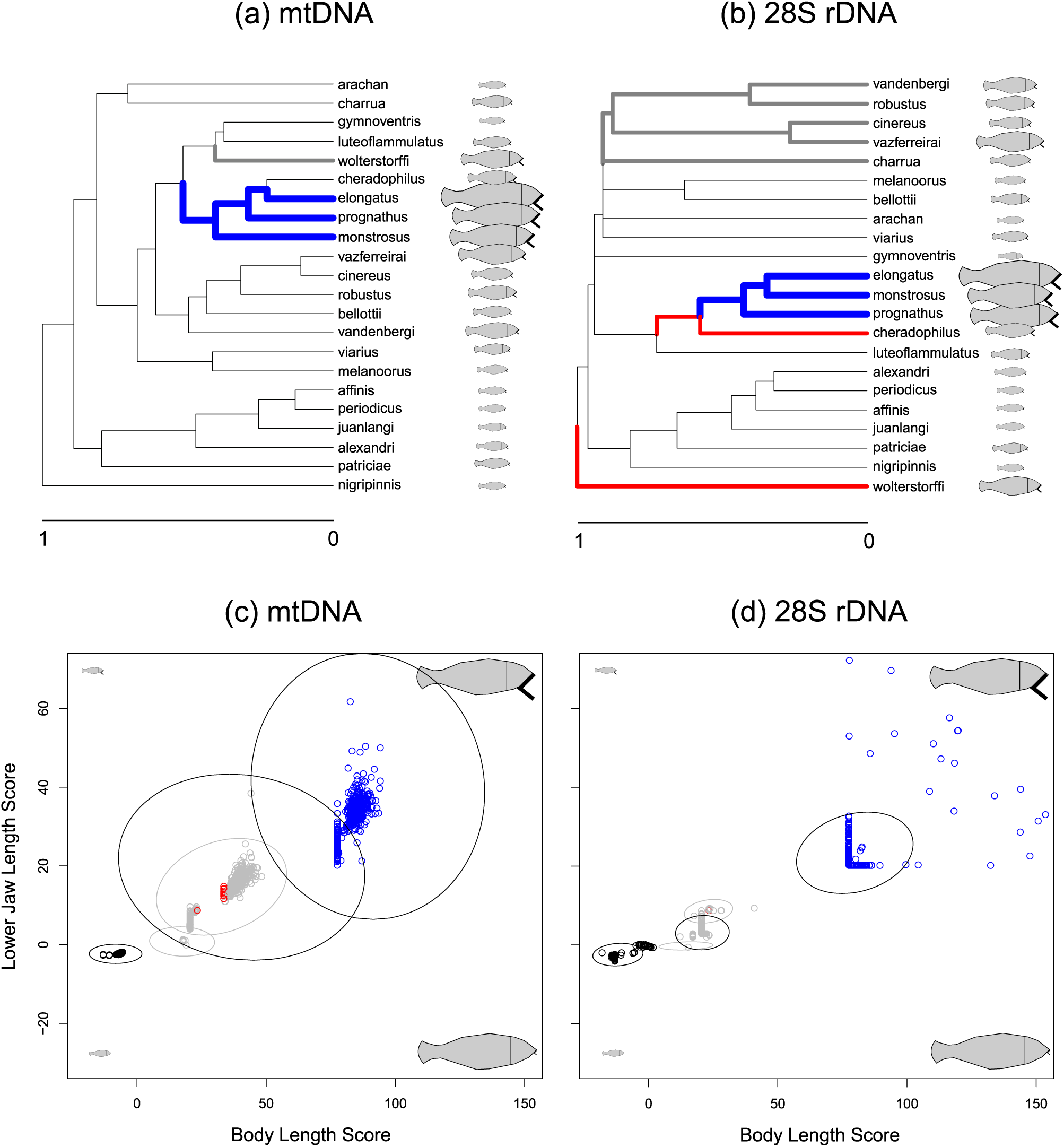
Top. Alternative consensus species trees for either (a) the three mitochondrial or (b) three 28S nuclear rDNA markers, with selection regimes of the selected OU model painted onto the branches. Panel (a) has three different selection optima and two regime shifts, (b) four selection optima and four shifts. Next to the trees, simplified fish representations show relative values of the size and jaw (black **<**) traits. Bottom. Scatter plots of estimated trait values at selection optima, where a model is selected and parameters estimated on each tree in a sample of 500 from the posterior distributions of species trees; (c) mtDNA data and (d) 28S rDNA data. Fish representations indicate changes of the size and jaw pc’s along the axes. The regimes with the smallest trait values at the selection optima are plotted as black points, the regimes with the largest trait values in each model with blue points. Intermediate selection regimes are in grey (when 3 regimes are selected) or red (when 4 regimes are selected). Per selection regime, ellipsoids are added containing 95% of the posterior density in parametric bootstrap simulations of the trait values at the selective optima. The ellipsoid contours are grey for the selection regimes with intermediate trait values when selected models have four regimes and black otherwise. The selected models contain three selection regimes for 95% of the mtDNA and 28S species trees, in 2% (mtDNA) and 5 % (28S rDNA) of the cases two regimes are preferred, and for the remainder of the trees four regimes. The three black ellipsoids therefore represent the models selected in the majority of cases.

The large polytomy involving the placement of 34 of the 72 *Austrolebias* individuals in the 28S consensus phylogenetic trees in Figs. 3 and S3 corresponds to short branch lengths in the separate trees of the posterior distribution. Contrary to the results of Ponzetto *et al* (2015), we found *Ophthalmolebias* as a basal split in the mtDNA tree, and therefore rooted the nuclear tree on *O. suzarti*. Our two gene trees confirm that *A*. *apaii* and *bellottii* are one species, as already suggested by the results of Garcia *et al* (2012). The large C*ynolebias albipunctatus* apparently originated within *Hypsolebias* (Fig. S3) and is not a sister taxon of *Austrolebias*.

We observed discordance between nuclear and mitochondrial trees (Figs. 3, S3 and S6). This was the case for *S. chacoensis*, *A. nigripinnis* and for the placement of *A. cheradophilus* within one clade of large *Austrolebias* (Fig. 2). More importantly, some taxa with discordances show a geographic association: *A. wolterstorffi, gymnoventris* and *luteoflammulatus* are sister species in the mitochondrial tree and co-occur in The Patos Lagoon system and associated plains. In the nuclear 28S tree *A*. *wolterstorffi* is placed at the root of *Austrolebias* and *A. luteoflammulatus* is the sister taxon of the group of large species that includes *A*. *elongatus*. *Austrolebias periodicus*, *juanlangi* and *affinis* are grouped together in the mitochondrial tree and all three occur in the “Negro” area of endemism. In the 28S tree *A. affinis* is the sister taxon of *A. alexandri*, occurring in the “La Plata” area of endemism. These last two sets of species and their geographic associations suggest that introgression or hybridization might have taken place within each set (Toews and Brelsford 2012) and the true underlying phylogeny might be a reticulate network.

### Species Trees

We determined species trees for the genus *Austrolebias* using the posterior distribution for the mtDNA tree with full data on all individuals included (Fig. S6). Removing individuals with partial mtDNA data improved the properties of the consensus tree considerably. We used the posterior 28S nuclear trees that include all individuals with data (Fig. S3).

### Morphological Traits

Fitting multivariate OU models to the size and jaw scores and the mitochondrial trees resulted in 95% of the cases in a model with three selection regimes having the lowest AICc (Fig. 3a&c). In comparison to an OU model with a single optimum, adding selection regime shifts decreased the AICc on average with 45 (s.d. 1.7). These AICc changes across the posterior distribution of mtDNA trees corresponded to 0.0006 (s.d. 0.002, BM true model) and 0.0008 (s.d. 0.003, OU true model) average tail probabilities across the posterior distribution of trees. Hence BM and single regime OU models can be rejected. Note that 5% quantile AICc differences were on average -22 in the simulations, confirming that simulations of null hypotheses are crucial. When optimal trait values of different regime shifts were allowed to be equal, the AICc could be decreased further with an average of 9.6 (2.5).

Multivariate OU models fitted to the size and jaw scores and the nuclear phylogenetic trees most often selected a model with three selection regimes (95%, Fig. 3b&d). Adding selection regime shifts with selection to different optima decreased the AICc with 42.6 (4.2). These AICc changes corresponded to 0.0006 (s.d. 0.002, BM true model) and 0.0007 (s.d. 0.003, OU true model) average tail probabilities. BM and single regime OU models can thus be rejected again. Average five percent quantile AICc values were -19 (BM) and -22 (OU). When optimal trait values of different regime shifts were allowed to be equal, the AICc decreased a further 5.6 (1.4) on average.

In all selected models, optimal trait values for the smallest species are well separated from the others (Figs. 3c&d). The less resolved 28S trees classify intermediate optima as clearly different (Fig. 3d). For the better resolved mtDNA trees, the largest and second largest optimal trait values have overlapping 95% posterior density regions (Fig. 3c). In particular jaw length scores for the intermediate optima are consistently overlapping with those for the smallest species. Adding measurement errors of gradually increasing magnitude to both traits resulted in two selection optima being more often preferred than three (Figure S7). For the 28S trees, this already occurred when errors with a small variance were added. The large *A. elongatus*, *monstrosus* and *prognathus* are generally selected towards a different optimum than all other species irrespective of measurement error. Small shifts in the estimated values of the selection optima depend more strongly on the trees sampled from the posteriors than on the magnitude of measurement error (Fig. S7). We conclude from this assessment of measurement error that it can blur the detection of some selection optima, not that measurement error in our data would generate additional spurious selection regimes.

From inspection of the posterior distributions of phylogenetic trees, it seems appropriate to assess the plausibility of giant-dwarf speciation at the nodes leading towards the most recent common ancestor (*mrca*) of two clades with large species (*A. elongatus, prognathus, monstrosus, cheradophilus*; *A. vazferreirai, cinereus, robustus*) and at the node leading to *A. wolterstorffi*.

Changes at the mrca of *A. elongatus, prognathus* and *monstrosus* need to be assessed as well, as the 28S model for jaw length suggests that specialized piscivory evolved there. Table 1 lists how often regime shifts were fitted at these nodes, and how much selection optima were shifted. For the mtDNA trees, the most recent common ancestor of the clade containing *A. elongatus* shows a regime shift with changes in size and jaw length in nearly all trees (Table 1). Regime shifts towards *A. wolterstorffi* were fitted in many trees, but with much smaller jaw length changes with large standard deviation. These shifts can be considered negligible. The 28S trees all have a regime shift with large changes in body size and jaw length towards the subclade containing *A. elongatus, prognathus* and *monstrosus*. In half of the trees this shift is preceded by a different one involving *A. cheradophilus* and mainly for body length. We thus recover an element of the *gradual allopatric* scenario, with evolution of more specialized piscivory within a clade of large *Austrolebias*. Regime shifts towards *A. wolterstorffi* are preferred for more than half of the 28S trees, again with moderate but this time non-negligible changes in jaw length. The clade with *A. robustus, vazferreirai* and *cinereus* often has a regime shift for body length and jaw length on the 28S tree, but not for the mtDNA trees which resolve these species well.

**Table 1.** Robustness of results across posterior distributions of phylogenetic trees. Each analysis was repeated on a sample of 500 trees from the posterior distributions. Per node for a particular most recent common ancestor (MRCA), the fraction of trees with a regime shift is given and when a regime shift was predicted at that node, the average changes in the selection regime for body length (size) and jaw length scores (Surface) are given. For cladogenetic events leading to that node, the average proportion of stochastic character mappings (BioGeoBears) with sympatric speciation is given and the average rank of the rate of evolution (stabletraits) averaged across the posterior. Between brackets: standard deviations. For the 28S trees, BiogeoBears often converged to a model with no sympatric speciation. For that reason, we give average proportions of mappings with sympatric speciation at particular nodes for two groups of models (italic): with non-zero weights of sympatric speciation / with zero weights.

| Gene tree | MRCA *elongatus*,<br>*monstrosus*,<br>*prognathus*,<br>*cheradophilus* | MRCA *elongatus*,<br>*monstrosus*,<br>*prognathus* | Ancestor *wolterstorffi* | MRCA *robustus*,<br>*vazferreirai*,<br>*cinereus* |
| --- | --- | --- | --- | --- |
| Surface |  |  |  |  |
| mtDNA | 0.99<br>Size score 93 (23)<br>Jaw score 33 (16) |  | 0.98<br>Size score 43 (26)<br>Jaw score 16 (12) | 0.17<br>Size score 31 (6)<br>Jaw score 9 (3) |
| 28S rDNA | 0.50<br>Size score 38 (17)<br>Jaw score 7 (6) | 1.00<br>Size score 77 (26)<br>Jaw score 21 (8) | 0.57<br>Size score 33 (5)<br>Jaw score 6 (2) | 0.63<br>Size score 34 (6)<br>Jaw score 6 (2) |
| Stabletraits |  |  |  |  |
| mtDNA | Size score 40 (1)<br>Jaw score 42 (0) |  | Size score 34 (3)<br>Jaw score 40 (2) | Size score 18 (2)<br>Jaw score 28 (2) |
| 28S rDNA | Size score 36 (2)<br>Jaw score 41 (0) | Size score 42 (0)<br>Jaw score 42 (0) | Size score 39 (1)<br>Jaw score 40 (1) | Size score 24 (6)<br>Jaw score 15 (10) |
| BioGeoBEARS |  |  |  |  |
| mtDNA | 0.61 |  | 0.97 | 0.82 |
| 28S rDNA | *0.75 / 0* | *0.5 / 0* | *0.49 / 0* | *0.18 / 0* |

Trait changes towards the clade containing *A. elongatus* systematically rank among the fastest for size and jaw length (Table 1 and Figure 4). The rate on the branch towards *A. wolterstorffi* shows rapid change - even on the 28S trees - for jaw length and less extreme rates for body length. Evolutionary rates leading towards the clade with *A. robustus* are never very fast, thus rejecting the *giant-dwarf* scenario for that clade. For *A. wolterstorffi*, the non-significant change in the selection optimum for jaw length on the mtDNA trees leads to rejection of the scenario, but this might be a consequence of lack of statistical power and jaw length did change at a relatively fast rate. For the *elongatus* clade there is no reason to reject the *giant-dwarf* scenario or the variant with two successive speciation events with large trait changes.

**Figure 4.**
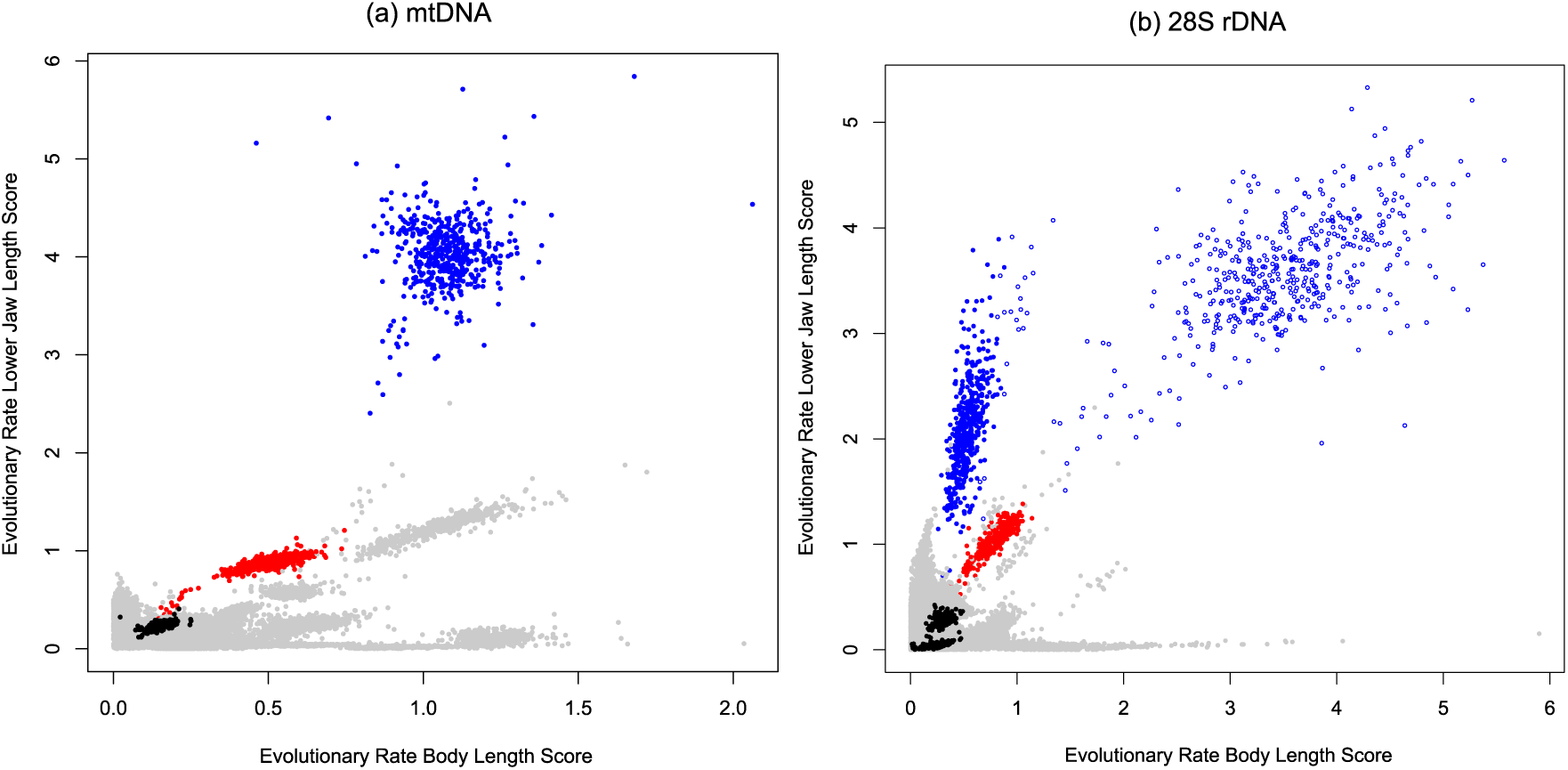
Rates of evolution on each branch in a phylogenetic tree estimated from a fit of stable trait distributions. Median rates for all branches in samples of 500 trees from the posterior tree distributions are represented, (a) for mtDNA species trees and (b) for 28S rDNA species trees. The rates on the branches leading towards *A. wolterstorffi* are in red, towards the most recent common ancestor of the three-species clade with *robustus* in black. The rates on the branches towards the most recent common ancestor of the four-species clade with *elongatus* are in blue. In the panel for the 28S trees, open blue points are added for the rates towards the most recent common ancestor of *elongatus, monstrosus* and *prognatus*.

### Biogeography

The rates of anagenetic dispersal and extinction in *Austrolebias* are consistently estimated to be zero. Model comparison between models with and without vicariance revealed that this model simplification reduced the AICc (mtDNA reduction 0.47 (s.d. 0.21); 28S rDNA 0.92 (s.d. 0.49)). We conclude that the *gradual allopatric* hypothesis is not supported. Allowing sympatric speciation decreases the AICc on average with 23.84 (1.7) across the mtDNA trees. For the nuclear 28S trees, it does not improve the model substantially. In 60% of the sampled trees, the probability of sympatric speciation is zero and the AICc of a model without sympatric speciation is on average 3.78 (6.1) smaller (Table 1). Since 40% of the 28S phylogenetic trees in the posterior do assign a non-zero probability to sympatric speciation, we decided to complete the analysis and not reject giant-dwarf speciation on the basis of average AICc alone.

Figure 5 shows ancestral ranges of largest marginal likelihood at each node in the consensus trees, for the full biogeographic model. The mtDNA tree has a single multi-area range, the La Plata – Negro range at the root of *Austrolebias* (Figure 5a). According to that reconstruction, area Negro has been colonized four times, La Plata and Western Paraguay each three times, and the Patos area of endemism once. Of the three events leading to large species, two occurred in Patos and one in La Plata. *Cynopoecilus* species immigrated in the Patos area and it can therefore not be excluded that these annual fish contributed to the evolution of piscivores. However, that should have happened in the presence of non-piscivorous *Austrolebias,* suggesting a second new scenario for the evolution of piscivory, where the arrival of a new potential prey triggers the evolution of piscivory from a non-piscivorous and small ancestor. Piscivory did not evolve in the Western Paraguay area and we do not need to consider the role of the annual killifish genera occurring there. Across the posterior tree distributions, most ancestral ranges are single-area ranges. Stochastic mapping placed a vicariance event at a node where large species evolved in less than 0.1% of the simulations, except for *A. wolterstorffi* in the 28S tree, where vicariance occurred in 49.5% of the simulations. We can reject the hypothesis that vicariance was involved in the origin of the large species of the *elongatus* and *robustus* clades.

**Figure 5.**
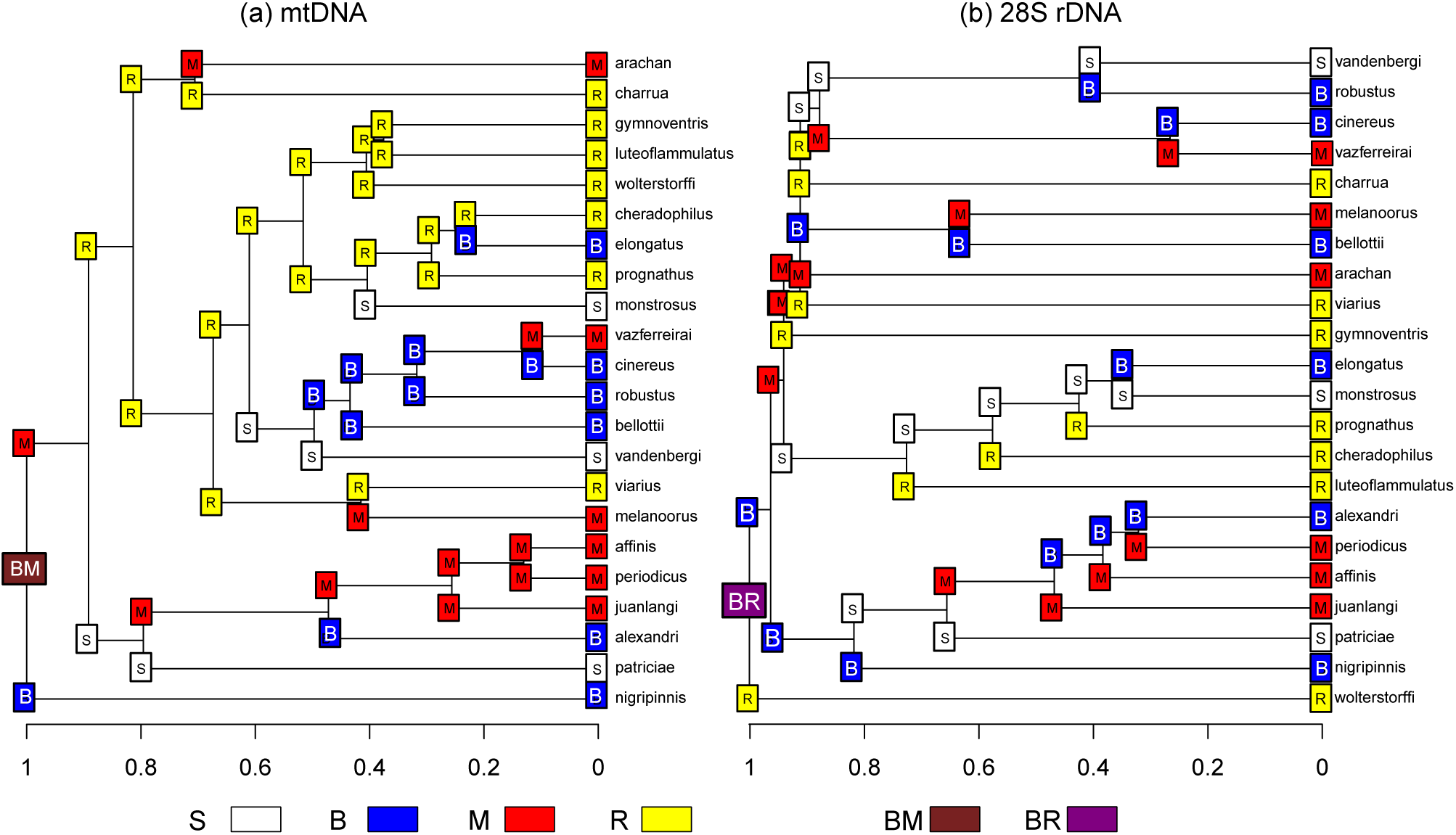
Biogeographical reconstruction of areas of endemism at ancestral nodes along consensus phylogenetic trees for (a) mtDNA (b) 28S rDNA data. Areas are abbreviated as indicated in Figure 2. B: the lower La Plata River basin and middle–lower Uruguay River basin, blue; R: Patos lagoon system and adjacent coastal plains to the south, yellow; S: the western part of the Paraguay River basin, white; M: Negro River drainage of the Uruguay River basin, and upper and middle sections of the Jacuí, Santa Maria, Jaguarão and Quaraí river drainages, red. BM and BR are ranges consisting of two areas of endemism. At each cladogenetic event in the tree, the descendant ranges with largest probability are shown.

We report additional results concerning the giant-dwarf scenario in Table 1. For the 28S trees, we present the results for trees in the posterior with a non-zero estimate of sympatric speciation (199/500) separately. When stochastic mapping was used to determine probabilities of event types given the mtDNA posterior tree distribution, tip ranges and model parameters, the split towards *A. wolterstorffi* had a probability of 0.97 to be a sympatric speciation. At the node towards the clade containing *A. vazferreirai* and *A. bellottii* this probability was 0.82. At the node towards the *elongatus* clade it was 0.61. When inspecting the models fitted to the 28S nuclear data that do not estimate a probability of sympatric speciation (Fig. 5b), it becomes clear that the predictions are not parsimonious, if jump dispersals are seen as events. In the clade with *A. elongatus* in the consensus tree, four dispersal events are favoured instead of two (Figure 5b).

For the clade containing *A. robustus* the overall probability that the split was a sympatric speciation is 0.07 given our 28S phylogenetic trees. Among the trees in the 28S posterior with non-zero weights for sympatric speciation, the probability of sympatric speciation is small for the clade containing *A. robustus*, intermediate for *A. wolterstorffi*, 0.75 for the split where large species diverge from *A. luteoflammulatus* and one half for the subsequent speciation event where *A. cheradophilus* originates.

When inspecting stochastic mapping results, it became clear that the probabilities of events differ between nodes directly connected to a tip and nodes two edges away from a tip (Figure 6). We inspected them separately to assess whether probabilities found at specific nodes were extreme.

**Figure 6.**
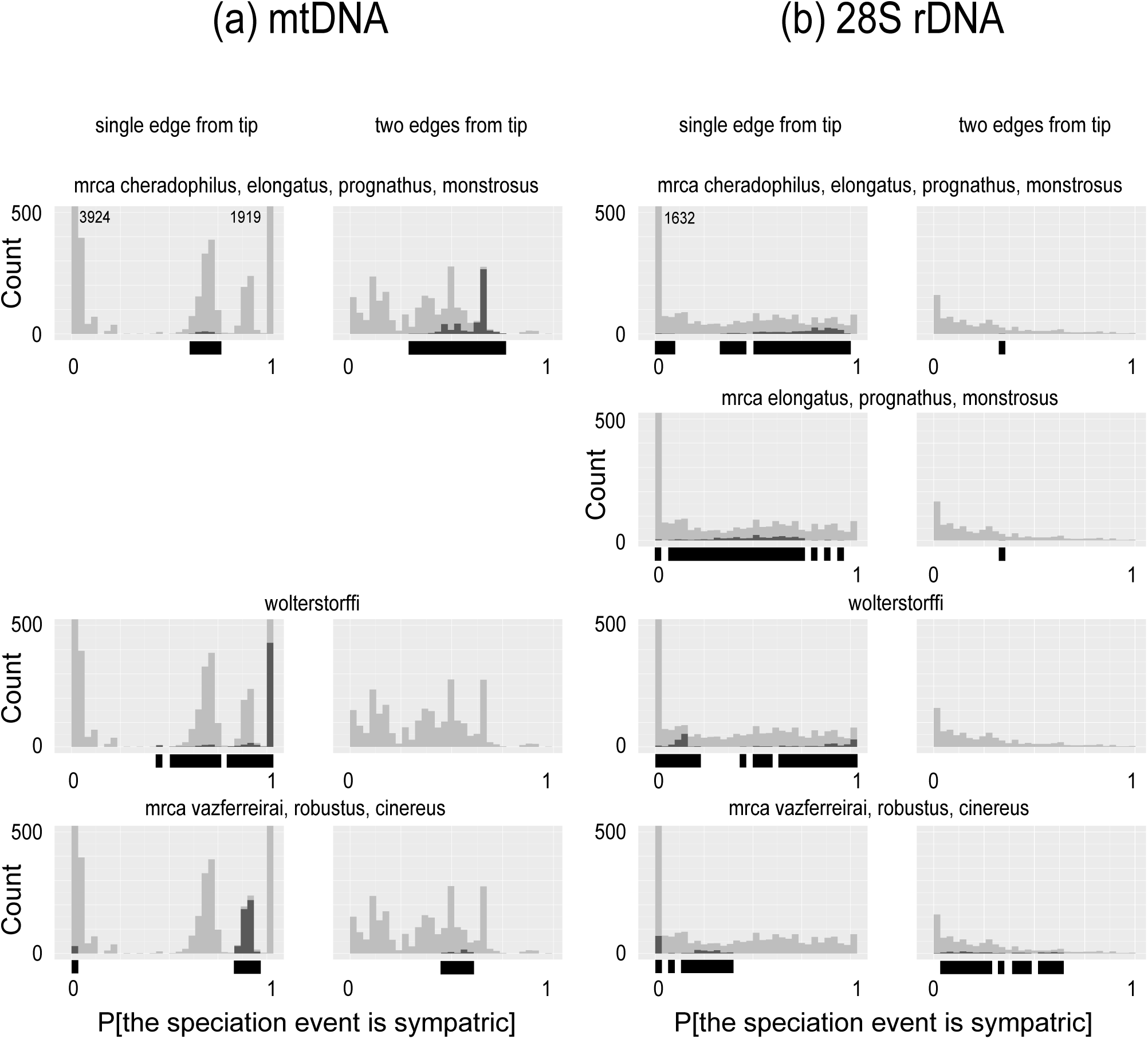
Results from stochastic mapping given samples of phylogenetic trees and selected biogeographic models. (a) Simulations on 500 trees sampled from the posterior for the mtDNA data, (b) 500 trees sampled from the posterior for the 28S rDNA data. Per node in a phylogenetic tree, the proportion of mappings that represent sympatric speciation is determined and histograms pooling these proportions over all trees and nodes are shown. We observed that these probabilities have very different distributions when nodes are directly connected to a tip or not. For that reason we plot these categories in separate columns. A count for a probability of one in a panel means that all 100 mappings for a node in a tree corresponded to sympatric speciation, a count for probability zero that no mappings for a node in a tree represented sympatric speciation. When extreme probabilities occurred much more often than intermediate ones, we adjusted the scale of the histogram to make these last ones easily visible and wrote the counts next to the bars for the extreme probabilities. The contributions of cladogenetic events at nodes immediately ancestral to the node written above the panel are colored black and superimposed on the pooled nodes in grey (mrca: most recent common ancestor). The events at these ancestral nodes are the ones we are most interested in. Under each histogram, the range of probabilities occurring for these nodes is indicated by a black bar. In column (b) we only represent the nodes of models that estimated a non-zero probability of sympatric speciation, i.e., which rejected the null hypothesis of no sympatric speciation.

We observe that simulated probabilities of sympatric speciation for each taxon with large species are consistently among the largest probabilities on the mtDNA trees, except for the clade with *robustus* (Fig. 6a). On the 28S rDNA trees and when a non-zero probability of sympatric speciation is estimated, the probability values per node are much more dispersed and smaller (Fig. 6b). This confirms that the 28S trees provide generally weaker support for the occurrence of sympatric speciation.

## Discussion

A scenario for the evolution of piscivory in *Austrolebias* with gradual evolution and vicariance within a single clade of large species (Costa 2010) can be rejected. The *Austrolebias* species with large body sizes are not forming a monophyletic group in either of the gene trees we constructed. The most specialized piscivores evolved from small species in one or two speciation events without vicariance and with significant selection regime shifts and the fastest rates of trait change.

Our results indicate that the mtDNA phylogenetic trees used so far to generate phylogenetic hypotheses for this genus (e.g. Garcia *et al* 2014) should be complemented with nuclear ones now that mito-nuclear discordance is observed. The species complexes described in Garcia *et al* (2014) are partially supported. Our analysis only retrieves the *elongatus* complex of large species with long jaws. *C. albipunctatus* is a sister taxon to some of the *Hypsolebias* species, and may have originated within a clade of *Hypsolebias* species. It is worthwhile to repeat the assessment carried out here for as many *Cynolebias* species as possible and investigate whether all speciation events in the genus were allopatric or not - and if not, whether giant-dwarf speciation could have occurred.

Our biogeographical analysis reconstructs ancestral ranges differently from Costa (2010), who hypothesized a three-area range at the root node of *Austrolebias.* For the mitochondrial species tree, a root range consisting of the La Plata – Negro areas is predicted. However, numerous small species were still missing from our analysis and including non-*Austrolebias* sister taxa in the reconstruction might lead to different inference on the geographic origin of the genus. Our nuclear 28S tree did not resolve all species sufficiently well and we should not interpret the predicted range at the root node. It is striking that the evolution of two clades with large species is predicted to have occurred in the region of the Patos Lagoons (mtDNA). Previous studies also hypothesized sympatric speciation between sister species in this region (Loureiro *et al* 2015). It has apparently been less affected by marine incursions in the Miocene (Hubert and Renno 2006). Combined with the lack of secondary immigration of *Austrolebias* species from elsewhere, this could have increased the scope for local diversification, even while *Cynopoecilus* immigrated.

The third taxon with large species (which includes *A. vazferreirai*) is more associated with the areas where marine incursions have taken place. The large sizes in these species may correlate with increased dispersal capabilities into newly available and more rapidly changing habitats.

### The plausibility of giant-dwarf speciation

Our analysis revealed three nodes where large species evolved and we assessed for each of these whether *giant-dwarf* speciation is a possible explanation. The clade containing *A. elongatus, prognathus* and *monstrosus* originated with rapid trait changes for body size and relative lower jaw length and the selection regime has shifted towards larger sizes and larger jaw lengths. Sympatric speciation is not rejected for this clade. We propose that the clade with *A*. *elongatus*, *monstrosus* and *prognathus* has originated by giant-dwarf speciation or by one of the two related scenarios suggested by our analysis. The first is suggested by the 28S trees where just body length showed evolutionary branching in a first event followed by body and jaw length divergence at a subsequent one. Theoretical modelling should investigate whether this scenario can occur in sympatry or requires allopatry/parapatry between the different large species. A second new scenario follows from inspecting our results in combination with the biogeographical analysis of the genus *Cynopoecilus* (Costa 2016). The temporary ponds of the Patos area where the largest jaw length and size divergences occurred, were colonised only once by *Austrolebias* (mtDNA) and by *Cynopoecilus*. Alternative to variation generated by within-species divergence, colonisations from elsewhere can generate trait divergence (Rundle and Schluter 2004).

Piscivorous species might have evolved after the immigration of *Cynopoecilus* into the Patos area. We note that this does not rule out sympatric speciation or that the small *Austrolebias* coexisting with large species and with *Cynopoecilus* became prey species.

For *A*. *wolterstorffi* the support for *giant-dwarf* speciation is weaker, but not weak enough to reject the hypothesis. Trait changes were smaller and slightly less rapid, but the species emerged with one of the fastest evolving jaw lengths. The placement of *A. wolterstorffi* in the mtDNA tree leads to a high predicted probability of sympatric speciation (speciation within an area of endemism), but this event might mark an introgression in sympatry, not a speciation event. The placement of this species near the root of the 28S tree lowers the probability that a trait shift is detected and renders the reconstructed ancestral range unreliable. We need to assess introgression in more detail for this species such that constraints on trait and range divergence can be determined more precisely. Costa’s (2010) analysis based on morphological characteristics placed this species among the species of the *elongatus* clade, while Costa (2009) identified tooth characteristics in this species indicative of molluscivory. Gut content data may show whether it is indeed such a specialist or whether piscivory does occur.

For the clade containing *A*. *vazferreirai*, *cinereus* and *robustus,* giant-dwarf speciation can be rejected. Rates of trait change are too low and there is no substantial jaw length evolution in the 28S trees. The probability of sympatric speciation for the node at the base of this clade is intermediate to low for a node connected to a descendant tip (mtDNA). The 28S trees nearly reject sympatric speciation for this clade.

Disruptive frequency dependent selection generally favours an increase in phenotypic variation. However, phenotypic variation can be increased in very different ways (Rueffler *et al* 2006) of which speciation is one, growth plasticity another (Claessen *et al* 2000) and genetic polymorphisms with cannibal morphs a third (Dercole and Rinaldi 2002). Why would *giant-dwarf* speciation have occurred and not one of the alternatives? Environmental cues explaining the divergence are not known and neither have genetic polymorphisms for size morphs been observed. The fact that all killifish hatch when rain fills the ponds combined with a large size divergence between incipient species may have resulted in assortative mating for size and thus generated a smaller probability of genetic exchange between size categories. Whatever the original mechanism that generated giants and dwarfs, the result would be speciation. On the other hand, species that are both under selection towards a shared size optimum might interbreed more easily and allow the introgressions suggested by the mito-nuclear discordances in our data.

We are aware that sympatric speciation remains a contentious issue and that the presence of sympatry strongly depends on the scale and detail of the ranges considered. Examples of *in situ* speciation have turned out to be more likely representing parapatric processes. In two examples of potential sympatric speciation in lacustrine fish (Schliewen *et al* 2001; Barluenga *et al* 2006), emerging species tend to specialize on different resources that can be unevenly distributed across habitats within a lake. The fact that piscivores and their prey need to meet, naturally limits the scope for parapatry and allopatry in giant-dwarf speciation. Such side conditions can make it easier to demonstrate sympatric speciation (Bolnick and Fitzpatrick 2007). However, the presence of other annual fish genera such as *Cynopoecilus* might increase the scope for parapatry among *Austrolebias* species again as they provide another source of prey for incipient piscivores.

### Limitations and prospects

We carried out a small additional investigation of the sensitivity of results in our biogeographical analysis to the presence or absence of species in our phylogenetic trees. When we pruned the mtDNA trees to remove four species where secondary contact might have been mistaken for sympatric speciation (*A*. *wolterstorffi*, *gynmoventris*, *periodicus*, *affinis*), zero probabilities for sympatric speciation were consistently estimated. If range changes and dispersal jumps are counted as events and range copying with sympatric speciation not, then unparsimonious reconstructions were made here with more events than necessary (Fig. 5b). When all pairs of sister species included in the tree occupy pairwise different ranges, the probability of sympatric speciation seems to be forced towards zero (Ree and Sanmartin 2018). It thus seems likely that this type of analysis is sensitive to the range patterns of missing species, as predictions at deeper nodes can strongly depend on the occurrence of events near tip nodes. Fortunately, stratification of this type of analysis is already possible (Matzke 2014), with different parameter values in different parts of a tree. However, whether stratification can remedy issues caused by missing data is unclear. Generally, the noise generated by errors, phylogenetic uncertainty and missing species might bias the odds more against scenarios such as sympatric speciation requiring a very specific configuration of ranges in ancestral and descendant species.

Our analysis and results note that introgression between sympatric species could lead to overestimating the frequency of sympatric speciation on mtDNA phylogenetic trees, while incomplete lineage sorting in nuclear genes might blur geographic associations between species above effects of noise in the data and lead to underestimation of sympatric speciation. Our results based on the 28S gene trees suggest such underestimation, while the hints of introgression in the results suggest overestimation on the mtDNA tree. The true probability of sympatric speciation might then be in between the parameter estimates obtained from mtDNA and nuclear DNA trees.

At several instances in our analysis, we had to scrutinize results and reconsider data and models. We believe that this effort has paid off, given the limitations of the data used. Our data suggest that the true underlying phylogeny for *Austrolebias* might be a reticulate network. We based our analysis on a pair of trees that we interpreted as alternative species bifurcation histories both compatible with the network structure. This approach does not a priori exclude introgression, while it does not require a reticulate network to be estimated either. Phylogenetic comparative modelling now does start allowing for exchange between branches, by modelling hybridization effects on traits (Jhwueng and O’Meara 2015, Solís-Lemus *et al* 2017) but the models will inevitably become more parameter-rich hence sacrificing statistical power.

Measurement error in the morphological species traits could not be estimated jointly with multi-modal OU phylogenetic comparative models, motivating our simulation of measurement error effects. In future studies, replacing field measurements of phenotypic traits by lab measurements in standardized environments might allow for a more precise assessment of variability in species traits. The areas of endemism we used to delineate species ranges might be too crude as well, and more detailed range assessments would be useful. This study also motivates more ecological life history assessments. In the field, annual killifish occur in different local assemblages with the large species sometimes absent, which might facilitate investigations of current selective pressures on body size and jaw length. The demonstration of multi-peaked adaptive landscapes or local fitness minima in the absence of piscivores would lend further indirect support to the plausibility of giant-dwarf speciation. However, since the main traits involved will be size-related, we will need to discriminate between this scenario and other types of character displacement (Schluter 2000; Pfennig and Pfennig 2010).

Given that our phylogenetic hypotheses were based on limited data, much scope seems left to reject our standing hypotheses with phylogenetic histories based on several nuclear genes. It will be useful to investigate whether introgression, hybridization and the associated genomic consequences are common in *Austrolebias*, also from a life history perspective. Hybridization events might be an alternative avenue towards diversification, leading to sudden jumps in trait values for a lineage.

## Conclusion

We reject the hypothesis that large piscivorous *Austrolebias* annual killifish evolved in a series of vicariance events with gradual evolution. We have demonstrated selection regime shifts for size and jaw length in the annual killifish genus *Austrolebias*, and estimated a non-zero probability of sympatric speciation (speciation within an area of endemism) for the genus. We have identified a particular speciation event (the node in the phylogeny where the clade containing *A. elongatus* originates) where, given all limitations of the methods and data used, giant-dwarf speciation cannot be rejected. This speciation scenario might need to be considered for other taxa as well.

## Acknowledgements

We thank Leonie Doorduin and Azahara Barra who collected part of the DNA sequence data while the first three authors worked at the Institute of Biology in Leiden. TVD was supported by an NWO Veni grant, the Amsterdam Treub foundation and received financial support from Killidata.org. VS and AJH were supported by NERC. Heber Salvia, Martin Fourcade, Lucho Lobo and Fabiana Cancino are thanked for help with obtaining samples.

## References

Barluenga M, Stoelting KN, Salzburger W et al (2006) Sympatric speciation in Nicaraguan crater lake cichlid fish. Nature 439:719–23

Bergsten J (2005) A review of long-branch attraction. Cladistics 21:163–193

Bernhart SH, Hofacker IL, Will S et al (2008) RNAalifold: improved consensus structure prediction for RNA alignments. BMC bioinformatics 9:474

Bolnick DI, Fitzpatrick B (2007) Sympatric speciation: theory and empirical data. Annu Rev Ecol Syst 38:459–487

Butler MA, King AA (2004) Phylogenetic comparative analysis: a modelling approach for adaptive evolution. Am Nat 164:683–695

Claessen D, de Roos AM, Persson L (2000) Dwarfs and giants: Cannibalism and competition in size-structured populations. Am Nat 155:219–237

Cook JR, Stefanski LA (1994) Simulation-extrapolation estimation in parametric measurement error models. J Am Stat Assoc 89:1314–1328

Costa WJEM (2003) Three new annual fishes of the genus *Aphyolebias* Costa, 1998 (Cyprinodontiformes: Rivulidae) from Bolivian and Peruvian Amazon. Comunicacoes do Museu de Ciencias e Tecnologia da PUCRS Serie Zoologia 16:155–166

Costa WJEM (2006) The South American annual killifish genus *Austrolebias* (Teleostei: Cyprinodontiformes: Rivulidae): phylogenetic relationships, descriptive morphology and taxonomic revision. Zootaxa 1213:1–162

Costa WJEM (2009) Trophic radiation in the South American annual killifish genus *Austrolebias* (Cyprinodontiformes: Rivulidae). Ichthyol Explor Freshwaters 20:179–191

Costa WJEM (2010) Historical biogeography of cynolebiasine annual killifishes inferred from dispersal–vicariance analysis. J Biogeogr 37:1995–2004

Costa WJEM (2011) Parallel evolution in ichthyophagous annual killifishes of South America and Africa. Cybium 35:39–46

Costa WJEM (2014) *Austrolebias araucarianus*, a new seasonal killifish from the Iguaçu river drainage, southern Brazilian Araucarian Plateau Forest (Cyprinodontiformes: Rivulidae). Ichthyol Explor Freshwaters 25:97–101

Costa WJEM (2016) Inferring evolution of habitat usage and body size in endangered, seasonal Cynopoeciline killifishes from the South American Atlantic forest through an integrative approach (Cyprinodontiformes: Rivulidae). PloS one 11:e0159315

Costa WJEM, Lacerda MTC, Tanizaki K (1988) Description d’une nouvelle espece de Cynolebias des plaines cotieres du Bresil sud-oriental (Cyprinodontiformes, Rivulidae). Revue Francaise d’Aquariologie Herpetologie 15:21–24

Costa WJEM, Amorim PF, Mattos JLO (2018) Synchronic historical patterns of species diversification in seasonal aplocheiloid killifishes of the semi-arid Brazilian Caatinga. PLoS ONE 13:e0193021

Dercole F, Rinaldi S (2002) Evolution of cannibalistic traits: scenarios derived from adaptive dynamics. Theor Popul Biol 62:365–74

Dorn A, Ng’oma E, Janko K et al (2011) Phylogeny, genetic variability and colour polymorphism of an emerging animal model: the short-lived annual *Nothobranchius* fishes from southern Mozambique. Mol Phylogenet Evol 61:739–749

Dorn A, Musilová Z, Platzer M et al (2014). The strange case of East African annual fishes: aridification correlates with diversification for a savannah aquatic group? BMC evol biol 14:1

Elliot MG, Mooers AØ (2014) Inferring ancestral states without assuming neutrality or gradualism using a stable model of continuous character evolution. BMC Evol Biol 14:226

García G, Wlasiuk, G, Lessa EP (2000) High levels of mitochondrial cytochrome b divergence and phylogenetic relationships in the annual killifishes of the genus *Cynolebias* (Cyprinodontifomes, Rivulidae). Zool J Linn Soc 129:93–110

García G, Alvarez-Valin F, Gómez N (2002) Mitochondrial genes: signals and noise in phylogenetic reconstruction within killifish genus *Cynolebias* (Cyprinodontiformes, Rivulidae). Biol J Linn Soc Lond 76:49–59

García G, Gutiérrez V, Vergara J et al (2012) Patterns of population differentiation in annual killifishes from the Paraná–Uruguay–La Plata Basin: the role of vicariance and dispersal. J Biogeogr 39:1707–1719

García G, Gutiérrez V, Ríos N et al (2014.) Burst speciation processes and genomic expansion in the neotropical annual killifish genus *Austrolebias* (Cyprinodontiformes, Rivulidae). Genetica 142:87–98

Garman S (1895) The cyprinodonts. 19:1–179

Geritz SAH, Kisdi É, Meszéna G, Metz JAJ (1998) Evolutionarily singular strategies and the adaptive growth and branching of the evolutionary tree. Evol Ecol 12:35–57

Ho LST, Ané, C (2014) Intrinsic inference difficulties for trait evolution with Ornstein-Uhlenbeck models. Meth Ecol Evol 5:1133–1146

Hofacker IL, Fontana W, Stadler PF et al (1994) Fast folding and comparison of RNA secondary structures. Monatshefte f Chemie 125:167–188

Hubert N, Renno J-F (2006) Historical biogeography of South American freshwater fishes. J Biogeogr 33:1414–1436

Huelsenbeck JP, Ronquist F (2001) MRBAYES: Bayesian inference of phylogeny. Bioinformatics 17:754–755

Ingram T, Mahler DL (2014) SURFACE: detecting convergent evolution from comparative data by fitting Ornstein-Uhlenbeck models with stepwise Akaike Information Criterion. Meth Ecol Evol 4:416–425

Jeanmougin F, Thompson JD, Gouy M et al (1998) Multiple sequence alignment with Clustalx. Trends Biochem Sci 23:403–405

Jhwueng D-C, O’Meara B (2015) Trait evolution on phylogenetic networks. biorxiv.org/content/early/2015/08/05/023986

Khabbazian, M, Kriebel, R, Rohe, K, Ané, C (2016). Fast and accurate detection of evolutionary shifts in Ornstein-Uhlenbeck models. Meth Ecol Evol 7:811–824

Liu L, Pearl DK (2007) Species trees from gene trees: Reconstructing Bayesian posterior distributions of a species phylogeny using estimated gene tree distributions. Syst Biol 56:504–514

Liu L, Yu L, Pearl DK (2010) Maximum tree: a consistent estimator of the species tree. J Math Biol 60:95–106

Loureiro M, Borthagaray A, Hernández D et al (2015) Austrolebias in space: scaling from ponds to biogeographical regions. In: Berois N, García G. de Sá RO, (eds) Annual Fishes: Life History Strategy, Diversity and Evolution. CRC Press, Boca Raton FL, pp 111–132

Losos JB, Glor RE (2003) Phylogenetic comparative methods and the geography of speciation. Trends Ecol Evol 18:220–227

Mallet J., Besansky N, Hahn MW (2015) How reticulated are species? BioEssays 38:140–149

Matzke NJ (2014) Model selection in historical biogeography reveals that founder-event speciation is a crucial process in island clades. Syst Biol 63:951–970

Mossel E, Roch S (2010) Incomplete lineage sorting: consistent phylogeny estimation from multiple loci. IEEE/ACM Trans Comput Biol Bioinform 7:166–171

Müller K (2005) SeqState - primer design and sequence statistics for phylogenetic DNA data sets. Appl Bioinformatics 4:65–69

Nosil P (2012) Ecological Speciation. Oxford University Press.

Oviedo S, Rovira M, Garcia G (2016) Genomic isolated regions: Linkage groups in parental and laboratory hybrids between *Austrolebias adloffi* species group. In: Berois N, García G, de Sá RO (eds) Annual fishes: life history strategy, diversity and evolution. CRC Press, Boca Raton FL, pp 295–307

Persson L, Wahlström E, Byström P (2000) Cannibalism and competition in Eurasian perch: population dynamics of an ontogenetic omnivore. Ecology 81:1058–1071

Pfennig DW, Pfennig KS (2010) Character displacement and the origins of diversity. Am Nat 176:S26–S44

Philippe H, Vienne DMD, Ranwez V, Roure B, Baurain D, Delsuc F (2017). Pitfalls in supermatrix phylogenomics. Eur J Taxon 283:1–25

Ponzetto JM, Britzke R, Nielsen DT et al (2016) Phylogenetic relationships of *Simpsonichthys* subgenera (Cyprinodontiformes, Rivulidae), including a proposal for a new genus. Zool Script 45:394–406

Ree RH, Moore BR, Webb CO, Donoghue MJ (2005) A likelihood framework for inferring the evolution of geographic range on phylogenetic trees. Evolution 59:2299–2311

Ree RH, Sanmartín I (2018) Conceptual and statistical problems with the DEC+ J model of founder-event speciation and its comparison with DEC via model selection. J Biogeogr in press.

Ree RH, Smith SA (2008) Maximum likelihood inference of geographic range evolution by dispersal, local extinction, and cladogenesis. Syst Biol 57:4–14

Rueffler C, Van Dooren TJM, Leimar O, Abrams PA (2006) Disruptive selection and then what? Trends Ecol Evol 21:238–245

Rundle HD, Schluter D (2004) Natural selection and ecological speciation in sticklebacks. In:Dieckmann U, Doebeli M, Metz JAJ, Tautz D (eds.) Adaptive Speciation (Cambridge University Press, Cambridge UK pp 192–209

Sanderson MJ (2002) Estimating absolute rates of molecular evolution and divergence times: a penalized likelihood approach. Mol Bio Evol 19:101–109

Schliewen U, Rassmann K, Markmann M et al (2001) Genetic and ecological divergence of a monophyletic cichlid species pair under fully sympatric conditions in Lake Ejagham, Cameroon. Mol Ecol 10:1471–88

Schluter D (2000) Ecological character displacement in adaptive radiation. Am Nat 156:S4–S16

Schöniger M, von Haeseler A (1994) A stochastic model and the evolution of autocorrelated DNA sequences. Mol Phyl Evol 3:240–247

Silvestro D, Kostikova A, Litsios G et al (2015) Measurement errors should always be incorporated in phylogenetic comparative analysis. Meth Ecol Evol 6:340–346

Solís-Lemus C, Bastide P, Ané C (2017). PhyloNetworks: a package for phylogenetic networks. Mol Biol Evol 34: 3292–3298

Toews DP, Brelsford A (2012) The biogeography of mitochondrial and nuclear discordance in animals. Mol Ecol 21:3907–3930

Thompson JD, Gibson TJ, Plewniak F et al (1997) The CLUSTAL_X windows interface: flexible strategies for multiple sequence alignment aided by quality analysis tools. Nucleic Acids Res 24:4876–4882

Uyeda JC, Caetano DS, Pennell MW (2015) Comparative analysis of principal components can be misleading. Syst Biol 64:677–689.

Wallis, G. P., Cameron-Christie, S. R., Kennedy, H. L., Palmer, G., Sanders, T. R., & Winter, D. J. (2017). Interspecific hybridization causes long-term phylogenetic discordance between nuclear and mitochondrial genomes in freshwater fishes. Mol Ecol 26:3116–3127

Xie W, Lewis PO, Fan Y et al (2011) Improving marginal likelihood estimation for Bayesian phylogenetic model selection. Syst Biol 60:150–160

