## Supplementary Materials for "Scope for sympatric giant-dwarf speciation driven by cannibalism in South-American annual killifish (*Austrolebias*)"

**Supplementary Material**

This file contains

A. Table S1 with sample information.
B. Tables S2 and S3 with markers and primers and PCR reactions used.
C. Additional figures S1-S6 to clarify the selection procedure of individuals to include for species tree construction. The figures show consensus phylogenetic trees that resulted from including all individuals with data in the analysis, or all individuals with data for either all mitochondrial or all nuclear markers.
D. An assessment of the effects of measurement error on OU modelling (Figure S7).

Table S1. List of samples used in the fitting of molecular phylogenetic trees. Location names refer to site names used by amateurs and others when exchanging fish or eggs. We kept site denominations as on the samples received. "NA" indicates missing values.

| | | | | Genbank accession numbers | | | | | |
|---|---|---|---|---|---|---|---|---|---|
| Genus | species | site | Individual code | cytb | 12S | 16S | 28S C1-C2 | 28S C6-C7 | 28S C12-D12pr |
| **To be added in accepted version** | | | | | | | | | |

Table S2. Markers and primers used in this study.

| Marker | Forward primer | | Reverse primer | | |
|---|---|---|---|---|---|
| 16S | 16S ar-L | 5'-CGC CTG TTT ATC AAA AAC AT-3' | 16S br-H | 5'-CCG GTC TGA ACT CAG ATC ACG T-3' | Kocher et al. 1989; Palumbi 1996 |
| 28S C1-C2 | 28S C1 | 5'- ACC CGC TGA ATT TAA GCA T -3' | 28S C2 | 5'- TGA ACT CTC TCT TCA AAG TTC TTT TC -3' | this study |
| 28S C6-C7 | 28S C6 | 5'- TCA CCT GCC GAA TCA ACT AGC -3' | 28S C7 | 5'- ACT ACC ACC AAG ATC TGC AC -3' | this study |
| 28S C12-D12pr | 28S C12 | 5'- TTA TGA CTG AAC GCC TCT AAG -3' | 28S D12pr | 5'- TGA CTT TCA ATA GAT CGC AG -3' | this study |
| 12S | L1091 | 5'-AAA AAG CTT CAA ACT GGG ATT AGA TAC CCC ACT AT- 3' | H1478 | 5'-TGA CTG CAG AGG GTG ACG GGC GGT GTG T -3' | Palumbi 1996 |
| cyt-b | CB3H | 5'GGC AAA TAG GAA (AG)TA TCA TTC 3' | Gludge L | 5' TGA CTT GAA (AG)AA CCA (CT)CG TTG 3' | Palumbi 1996 |

Kocher TD, ThomasWK, Meyer A, Edwards SV, Pääbo S, Villablanca FX, Wilson AC (1989) Dynamics of mitochondrial DNA evolution in animals: amplification and sequencing with conserved primers. *Proc Nat. Aca. Sc. USA* **86**:6196-6200

Palumbi SR (1996) Nucleic acids II: The polymerase chain reaction. In: Hillis DM, Moritz C, Mable BK (eds) *Molecular Systematics*. Sinauer Associates, Inc, pp 205-247

Table S3. PCR mix and cycling conditions. Components of the mix are given in µl per reaction; 2 µl of sample was added to a total of 25 µl reaction volume. Cycling conditions are given as time in seconds at temperature in degrees Celsius. For all markers, 36 PCR cycles were run.

| | cyt-b | 16S | 28S C1-C2 | 28S C6-C7 | 28S C12-D12 | 12S |
|---|---|---|---|---|---|---|
| 10X buffer 15mM MgCl2 | 2.5 | 2.5 | 2.5 | 2.5 | 2.5 | 2.5 |
| PCR grade water | 18.65 | 17.15 | 18.65 | 18.65 | 18.65 | 18.65 |
| dNTP's 2mM each | 2 | 2 | 2 | 2 | 2 | 2 |
| For Primer 5 µM | 0.8 | 0.8 | 0.8 | 0.8 | 0.8 | 0.8 |
| Rev Primer 5 µM | 0.8 | 0.8 | 0.8 | 0.8 | 0.8 | 0.8 |
| MgCl2 25mM | 0 | 1.5 | 0 | 0 | 0 | 0 |
| *Taq* polymerase 5U/µl | 0.25 | 0.25 | 0.25 | 0.25 | 0.25 | 0.25 |
| Initial denaturation | 360s at 93 C | 300s at 93 C | 360s at 94 C | 360s at 94 C | 360s at 94 C | 360s at 93 C |
| Denaturation | 60s at 93 C | 30s at 93 C | 60s at 94 C | 60s at 94 C | 60s at 94 C | 60s at 94 C |
| Annealing | 60s at 45 C | 10s at 52 C | 60s at 57 C | 60s at 55 C | 60s at 53 C | 60s at 52 C |
| Extension | 60s at 72 C | 60s at 72 C | 90s at 72 C | 90s at 72 C | 90s at 72 C | 90s at 72 C |
| Final extension | 360s at 72 C | 420s at 72 C | 360s at 72 C | 360s at 72 C | 360s at 72 C | 360s at 72 C C |

Figure S1. Nuclear 28S consensus phylogenetic tree, all individuals with (partial) data are included. Individuals are represented by the initial letter of their genus name, their species name, location of origin (if available) and sample label to facilitate retrieval from the supplementary table S1 (O: *Ophthalmolebias*, S: *Spectrolebias*, N: *Nematolebias*, C: *Cynolebias,* H: *Hypsolebias*, A: *Austrolebias*). We did not abbreviate genus names for the most distant outgroup species. Branch length magnitudes are indicated by a scale bar. In further modeling of these data, we excluded the data on individuals: A. nigripinnis Villa Soriano 9; A. affinis Durazno 23; H. magnificus 33 and A. alexandri San Javier 22. Non-*Austrolebias* individuals have a grey background. Species with individuals that were removed are indicated with boxes and connectors.

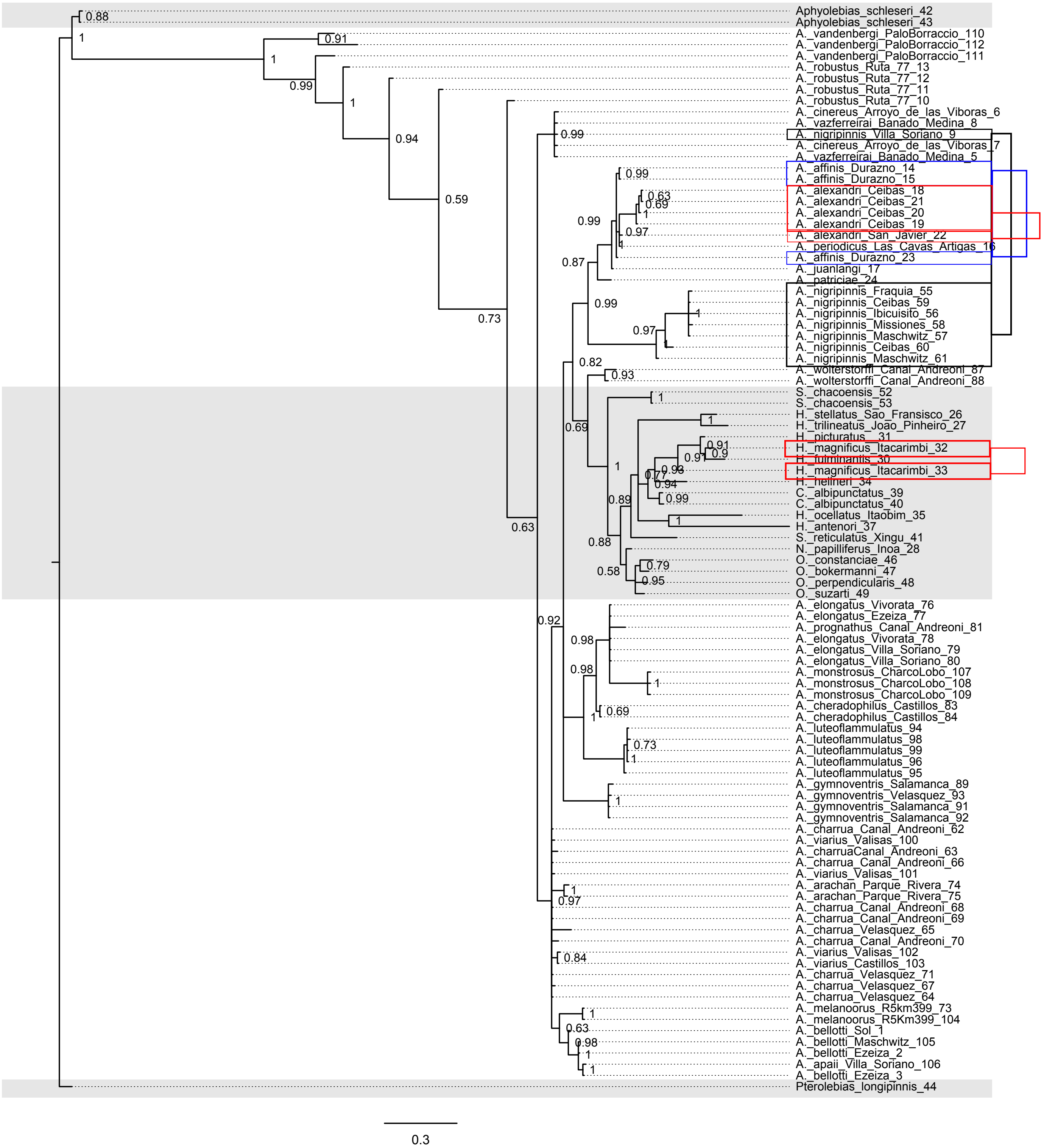

Figure S2. Nuclear 28S consensus tree, all individuals with full 28S data (three markers). In further modeling of these data, we excluded the data on individuals: A. affinis Durazno 23 and A. alexandri San Javier 22. Non-*Austrolebias* individuals have a grey background. Species with individuals that were removed are indicated with boxes and connectors.

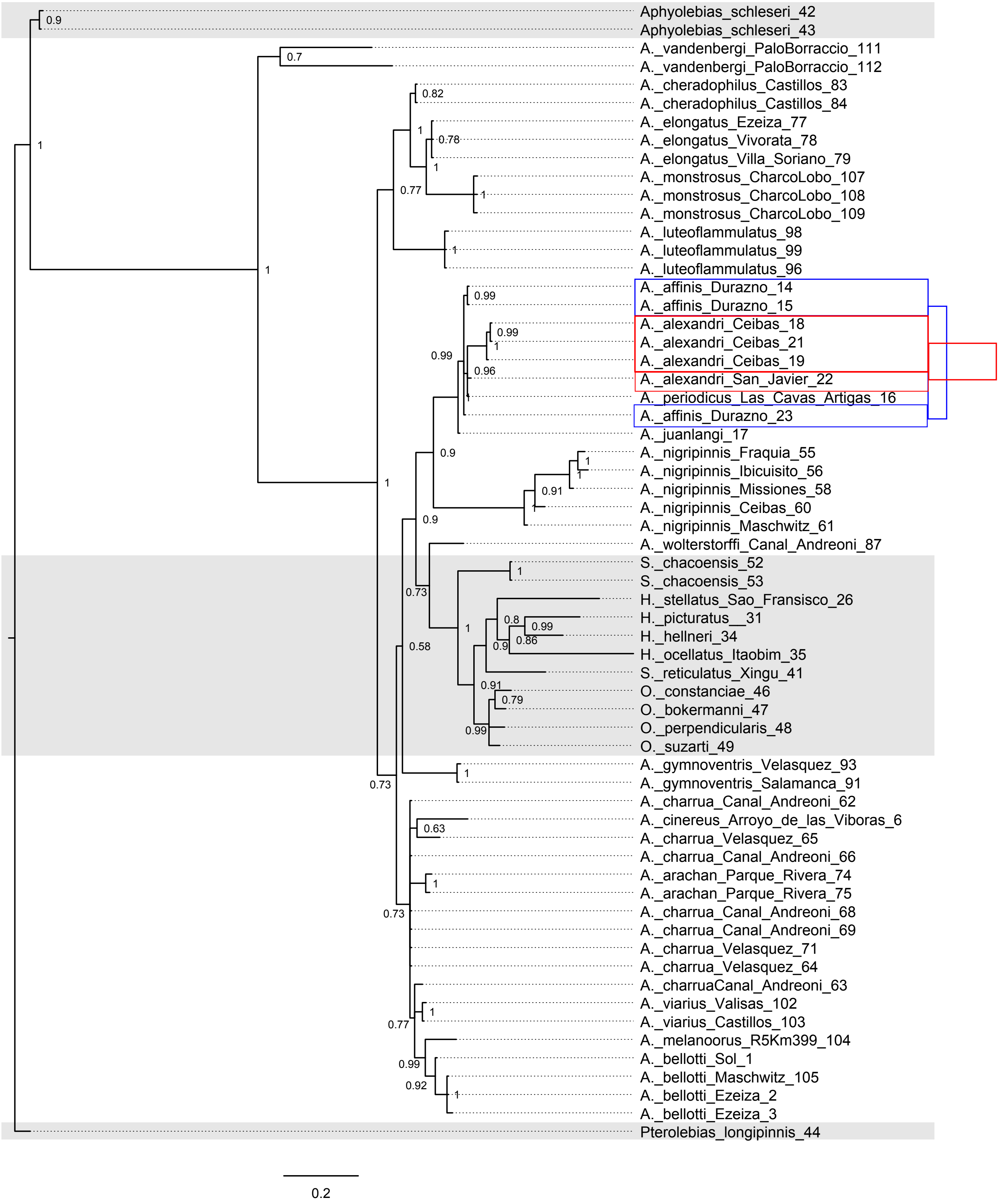

Figure S3. Nuclear 28S rDNA consensus tree of the posterior distribution used for trait modelling and biogeographic analysis. All individuals with data for at least one marker are included, except for a small number of individuals that did not group with kin or other individuals of their own species (see legend Figure S2), and outgroup individuals. Clades with large species are indicated by a blue background (*Cynolebias* and *Austrolebias*). Non-*Austrolebias* individuals by a grey background.

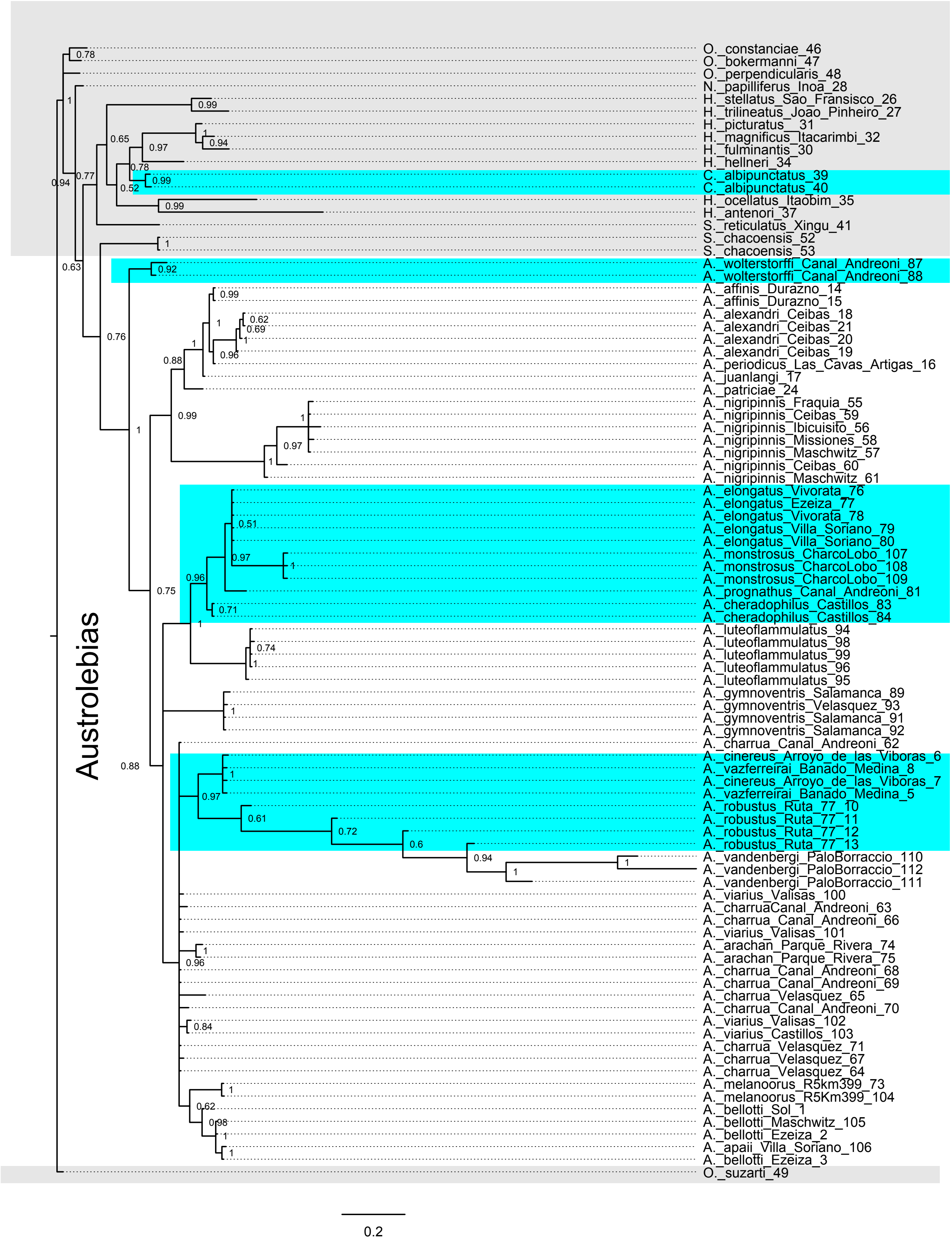

Figure S4. Mitochondrial consensus gene tree including all individuals with (partial) data. In further modelling of this dataset, we excluded the individuals: A. nigripinnis VS 9; A. alexandri San Javier 22; A. affinis Durazno 23;A.  luteoflammulatus 94; A. melanoorus 73 and A. cheradophilus La Paloma 86. Non-*Austrolebias* individuals have a grey background. Species with individuals that were removed are indicated with boxes and connectors.

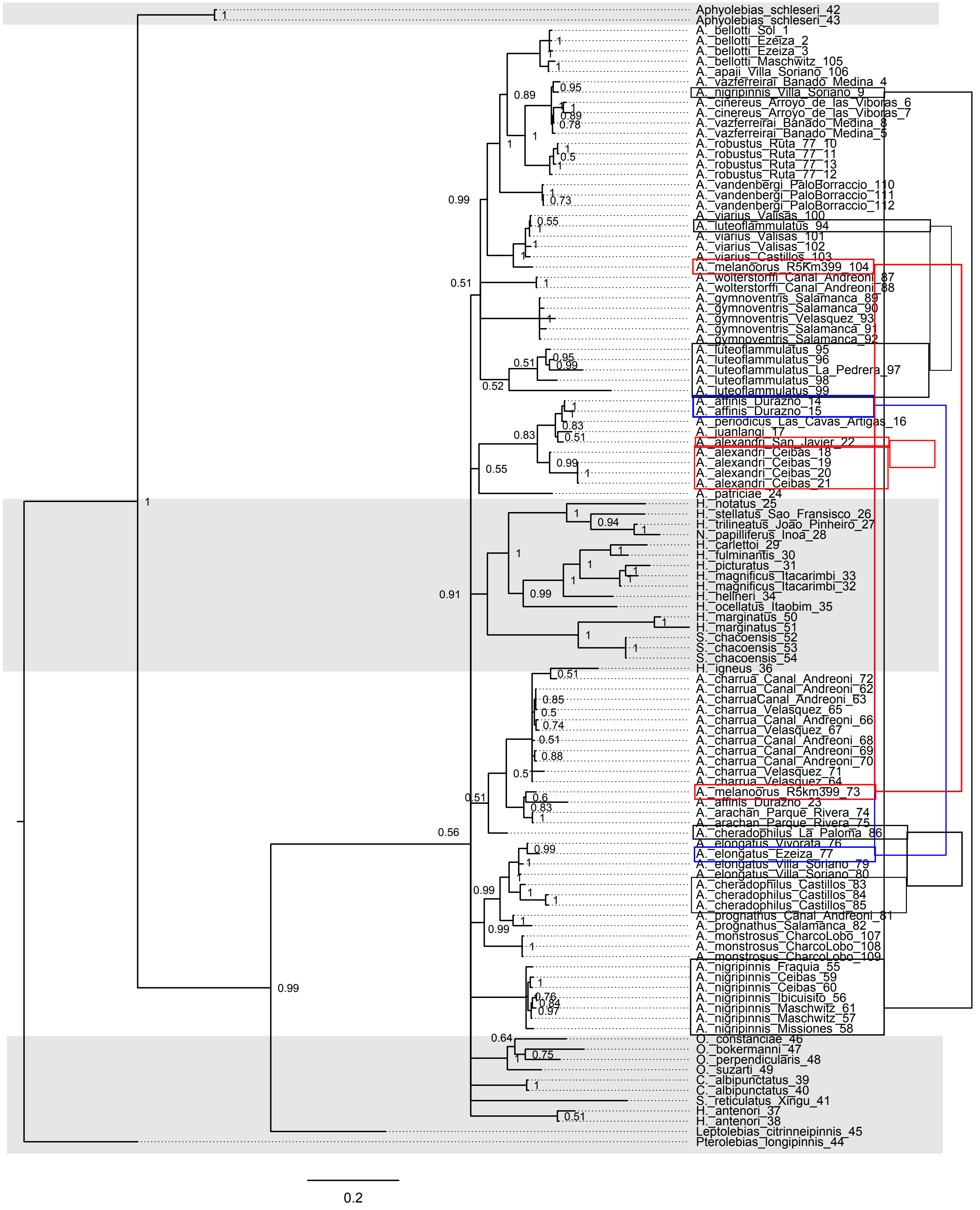

Figure S5. Mitochondrial consensus gene tree when only the individuals with data for the three markers are considered. In further modeling of these data, we excluded the data on individuals: *A. nigripinnis* Villa Soriano 9; *A. affinis* Durazno 23; *A. magnificus* 33; A. melanoorus 73 and *A. cheradophilus* La Paloma. Non-*Austrolebias* individuals have a grey background. Species with individuals that were removed are indicated with boxes and connectors.

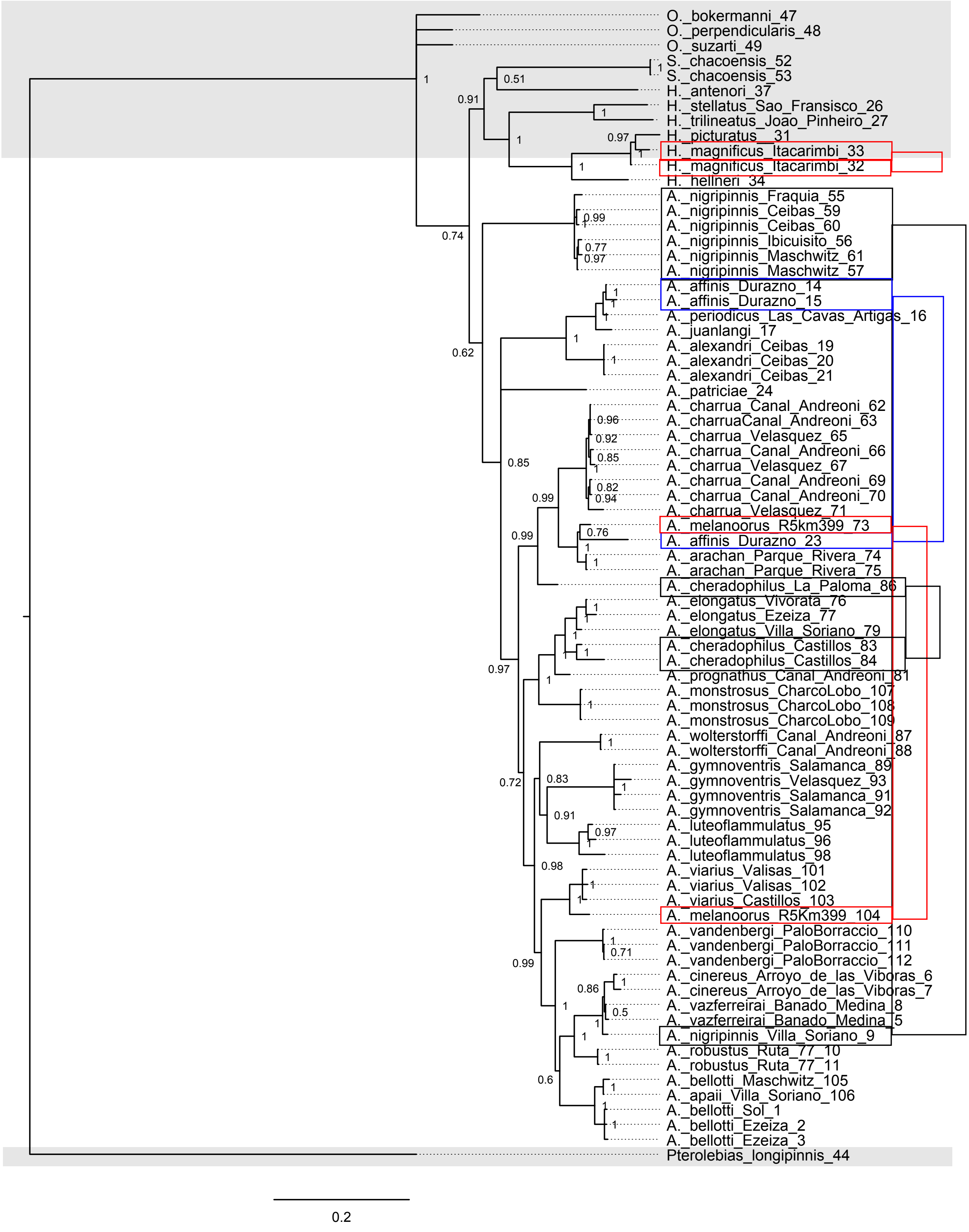

Figure S6. Mitochondrial consensus gene tree of the posterior distribution used for further analysis. All individuals with data for all three markers (cytb, 12S, 16S) are included, except for a small number of individuals that did not group with kin or other individuals of their own species (legend Figure S5). Clades with large species are indicated by a blue background (*Austrolebias*). Non-*Austrolebias* individuals by a grey background.

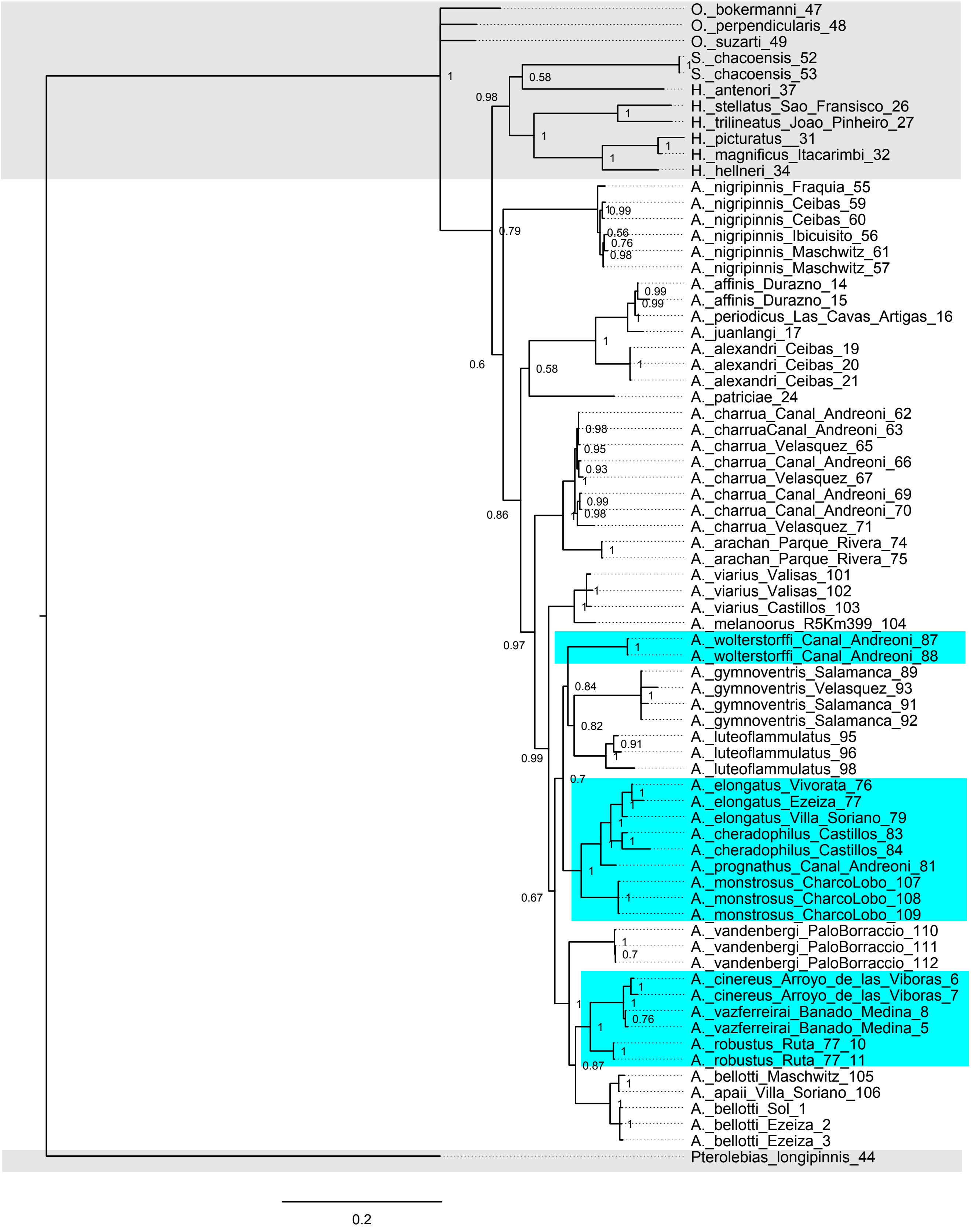

Figure S7. Assessment of effects of measurement error in the species traits on the estimation of optimal trait values for stabilizing selection and on the number of optima retained. This assessment used the posterior distributions of mitochondrial and 28S species trees and jaw length and body length scores as phenotypic species data. In each pseudo-dataset, independent Gaussian random variables with zero mean and a fixed standard deviation are added to both traits. Automatic OU model selection (Surface, Ingram & Mahler 2013) is then applied to this dataset and a random phylogenetic tree from the posterior. Pseudo-datasets were generated for standard deviations ranging from 0.05 to ten in 500 steps (x-axis). Colors indicate the number of selection optima retained per analysis: grey: two, black: three, red: four. Optimal trait values for body length and jaw length are plotted on the y-axis, all selection regimes per standard deviation.

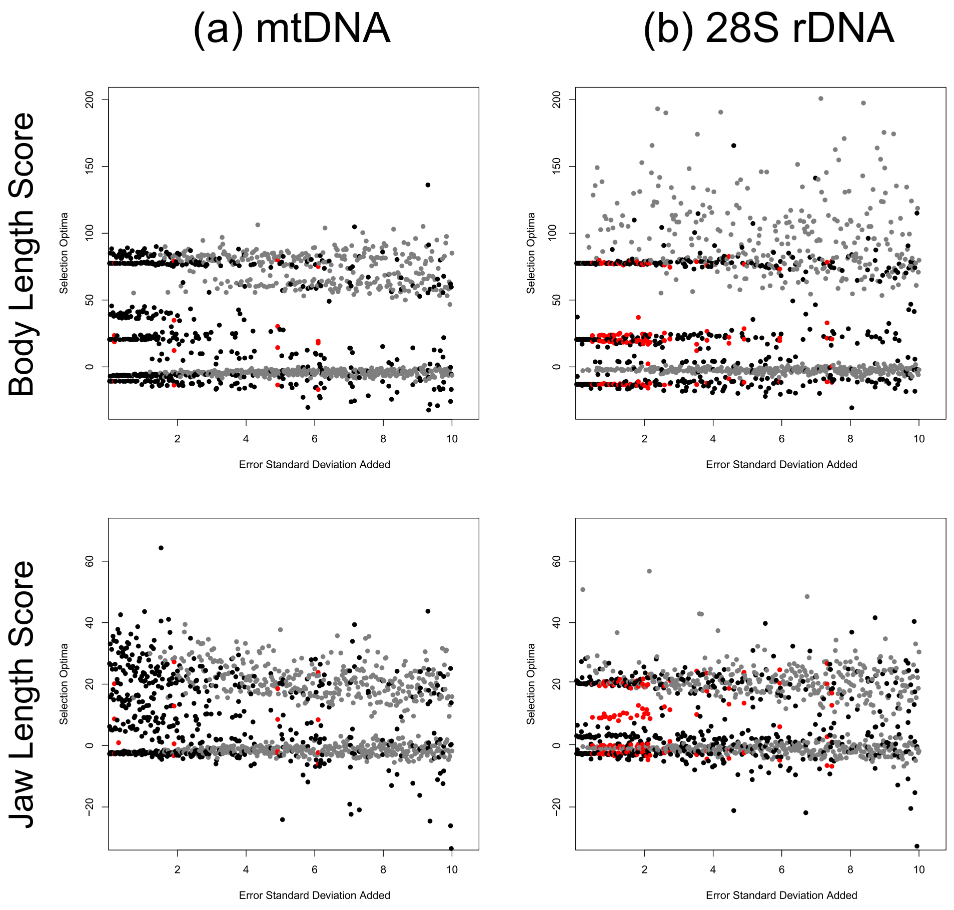
